## Supplemental Material for "Validity of fecal sampling for characterizing temporal variation in threespine stickleback’s gut microbiota"

Supplementary Figures

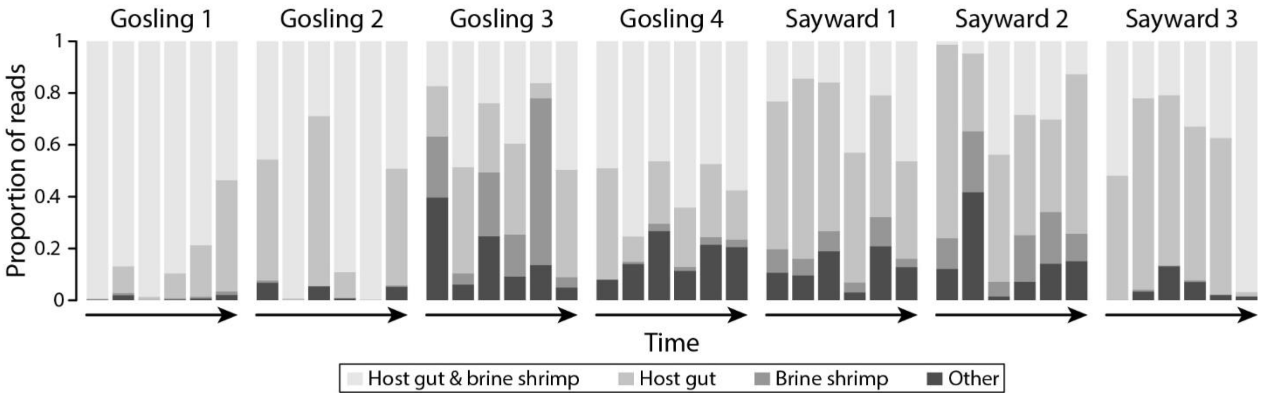


**Fig. S1:** Bar plots showing that large proportions (median: 90.3%) of most fecal bacterial communities were shared with the gut microbiota (host gut, host gut & brine shrimp). Smaller proportions were shared exclusively with diet (brine shrimp) or with neither the gut microbiota nor diet (other).

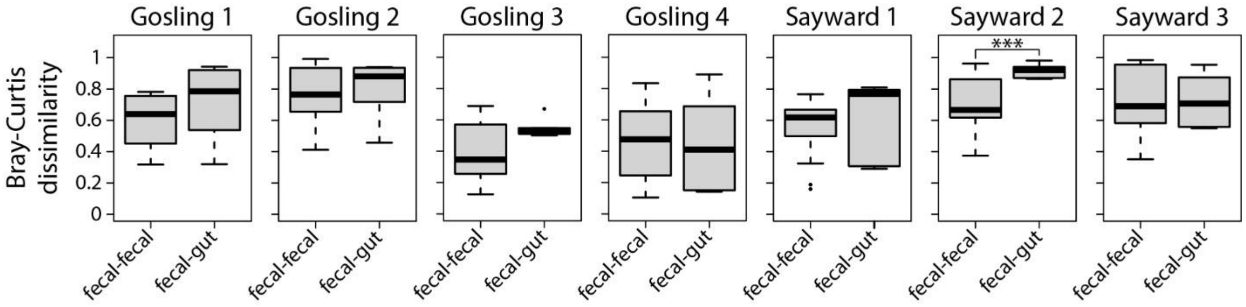


**Fig. S2:** Pairwise comparisons of Bray-Curtis dissimilarity revealed no significant differences among fecal samples and between fecal and gut samples for all but one individual (Sayward 2). Higher values indicate stronger dissimilarity among samples. Wilcoxon rank-sum tests, ***$P < 0.001$ (adjusted for multiple comparisons using Bonferroni correction).

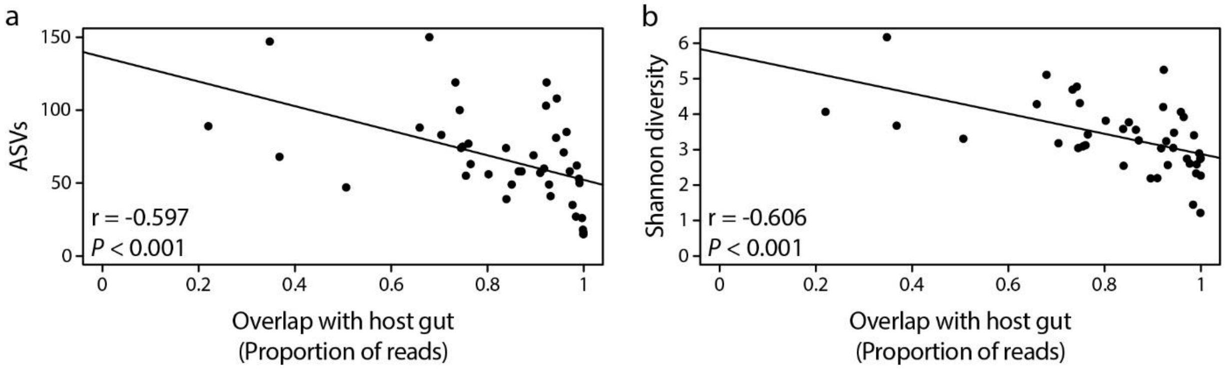


**Fig. S3:** Across fecal samples, both alpha diversity measures, the number of ASVs (a) and Shannon diversity (b), were negatively correlated with the overlap between bacterial communities of fecal samples and the gut tissue (based on Spearman's rank correlation coefficient).

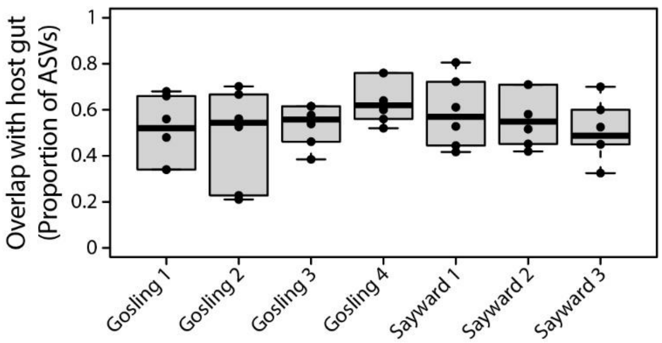


**Fig. S4:** The proportion of bacterial ASVs of the intestinal tissue that were also detected in fecal samples of the same individual differed substantially within and across individuals and ranged from 21.1 – 81% (median: 56%).

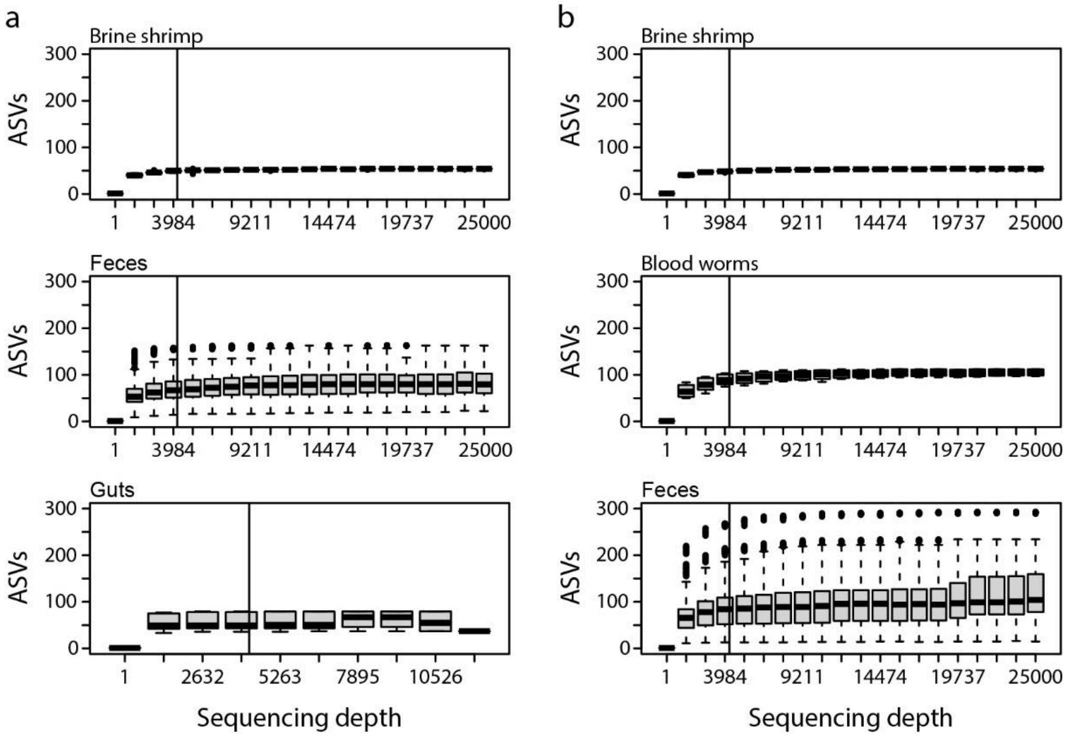


**Fig. S5:** Alpha diversity estimates (number of ASVs) at different rarefaction depths for different source materials (feces, guts, diet items) of the first (a) and second (b) experiment. The investigated sequencing depths range from 1 to 25,000 reads. We chose 4238 and 4275 reads as the sequencing depths at which the data was rarefied for the first and second experiment, respectively (indicated by vertical lines). At these sampling depths, a large proportion of the microbial diversity is captured across the different sample types.

**Table S1:** Test statistics of differential abundance analysis between fecal and gut samples in the first experiment using ANCOM in QIIME2 on the phylum level, with and without filtering of diet-derived bacteria. Test statistics of bacterial phyla that significantly differ between the two tissue types are indicated in bold.

| | no diet filtering | | diet filtering | |
|---|---|---|---|---|
| pyhlum | clr | W | clr | W |
| D_0__Bacteria;D_1__Deinococcus-Thermus | **-2.86243023** | **13** | **2.84700751** | **13** |
| D_0__Bacteria;D_1__Proteobacteria | 1.42329602 | 7 | -1.21216317 | 4 |
| D_0__Bacteria;D_1__Chlamydiae | 1.50827114 | 3 | -1.52369387 | 3 |
| D_0__Bacteria;D_1__Verrucomicrobia | -1.22065019 | 3 | 1.20522747 | 3 |
| D_0__Bacteria;D_1__Firmicutes | -1.06929815 | 2 | 1.05387543 | 1 |
| D_0__Bacteria;D_1__Nitrospirae | 0.71800218 | 2 | -0.73342491 | 2 |
| D_0__Bacteria;D_1__Spirochaetes | -0.4183669 | 2 | 0.40294418 | 2 |
| D_0__Bacteria;D_1__Gemmatimonadetes | 0.10966678 | 2 | -0.12508951 | 1 |
| D_0__Bacteria;D_1__Dependentiae | 0.00841432 | 2 | -0.02383705 | 1 |
| D_0__Bacteria;D_1__Bacteroidetes | 0.68617261 | 1 | -0.71223276 | 1 |
| D_0__Bacteria;D_1__Actinobacteria | 0.67476761 | 1 | -0.69019034 | 1 |
| D_0__Bacteria;D_1__Chloroflexi | 0.23580415 | 1 | -0.25122688 | 1 |
| D_0__Bacteria;D_1__Planctomycetes | 0.15682791 | 1 | -0.17225063 | 1 |
| D_0__Bacteria;D_1__Hydrogenedentes | 0.04952274 | 1 | -0.06494546 | 1 |

**Table S2:** Test statistics of differential abundance analysis between fecal and gut samples in the first experiment using ANCOM in QIIME2 on the genus level, with and without filtering of diet-derived bacteria. Test statistics of bacterial genera that significantly differ between the two tissue types are indicated in bold.

| | no diet filtering | | diet filtering | |
|---|---|---|---|---|
| genus | clr1 | W1 | clr | W |
| D_0__Bacteria;D_1__Proteobacteria;D_2__Gammaproteobacteria;D_3__Alteromonadales;D_4__Alteromonadaceae;D_5__Rheinheimera | **3.882167485** | **164** | NA | NA |
| D_0__Bacteria;D_1__Proteobacteria;D_2__Gammaproteobacteria;D_3__Betaproteobacteriales;D_4__Burkholderiaceae;D_5__Burkholderia-Caballeronia-Paraburkholderia | -2.543114732 | 146 | **2.511803739** | **144** |
| D_0__Bacteria;D_1__Proteobacteria;D_2__Gammaproteobacteria;D_3__Pseudomonadales;D_4__Moraxellaceae;D_5__Acinetobacter | -3.524947917 | 145 | **3.493636924** | **143** |
| D_0__Bacteria;D_1__Proteobacteria;D_2__Alphaproteobacteria;D_3__Rhodobacterales;D_4__Rhodobacteraceae;D_5__Gemmobacter | -3.477825377 | 141 | **3.446514384** | **139** |
| D_0__Bacteria;D_1__Deinococcus-Thermus;D_2__Deinococci;D_3__Thermales;D_4__Thermaceae;D_5__Meiothermus | -2.26990616 | 134 | **2.238595167** | **132** |
| D_0__Bacteria;D_1__Proteobacteria;D_2__Gammaproteobacteria;D_3__Betaproteobacteriales;D_4__Chitinibacteraceae;D_5__Chitinibacter | 3.550684957 | 120 | -3.581995949 | 121 |
| D_0__Bacteria;D_1__Proteobacteria;D_2__Gammaproteobacteria;D_3__Betaproteobacteriales;D_4__Burkholderiaceae;D_5__Schlegelella | -2.28263049 | 119 | 2.251319497 | 117 |
| D_0__Bacteria;D_1__Proteobacteria;D_2__Gammaproteobacteria;D_3__Aeromonadales;D_4__Aeromonadaceae;D_5__Aeromonas | 3.231030293 | 38 | -3.262341286 | 38 |
| D_0__Bacteria;D_1__Proteobacteria;D_2__Gammaproteobacteria;D_3__Oceanospirillales;D_4__Halomonadaceae;D_5__Halomonas | -2.096732272 | 30 | 2.065421279 | 29 |
| D_0__Bacteria;D_1__Deinococcus-Thermus;D_2__Deinococci;D_3__Thermales;D_4__Thermaceae;D_5__Thermus | -2.014231308 | 28 | 1.982920315 | 27 |
| D_0__Bacteria;D_1__Proteobacteria;D_2__Gammaproteobacteria;D_3__Alteromonadales;D_4__Shewanellaceae;D_5__Shewanella | 2.809573267 | 24 | -2.840884259 | 24 |
| D_0__Bacteria;D_1__Firmicutes;D_2__Bacilli;D_3__Bacillales;D_4__Bacillaceae;D_5__Geobacillus | -1.771525238 | 19 | 1.740214245 | 18 |
| D_0__Bacteria;D_1__Verrucomicrobia;D_2__Verrucomicrobiae;D_3__Verrucomicrobiales;D_4__Rubritaleaceae;D_5__Haloferula | -1.559007268 | 17 | 1.527696275 | 16 |
| D_0__Bacteria;D_1__Actinobacteria;D_2__Actinobacteria;D_3__Corynebacteriales;D_4__Mycobacteriaceae;D_5__Mycobacterium | 2.183095551 | 16 | -2.214406544 | 16 |
| D_0__Bacteria;D_1__Planctomycetes;D_2__OM190;Ambiguous_taxa;Ambiguous_taxa;Ambiguous_taxa | 1.557814442 | 13 | -1.589125435 | 13 |
| D_0__Bacteria;D_1__Actinobacteria;D_2__Actinobacteria;D_3__Micrococcales;D_4__Microbacteriaceae;__ | 1.97482149 | 11 | -2.006132482 | 11 |
| D_0__Bacteria;D_1__Proteobacteria;D_2__Alphaproteobacteria;D_3__Rhodobacterales;D_4__Rhodobacteraceae;__ | -2.490571568 | 11 | 2.459260575 | 10 |
| D_0__Bacteria;D_1__Bacteroidetes;D_2__Bacteroidia;D_3__Chitinophagales;D_4__Saprospiraceae;D_5__uncultured | 2.109323748 | 10 | -2.14063474 | 11 |
| D_0__Bacteria;D_1__Actinobacteria;D_2__Thermoleophilia;D_3__Solirubrobacterales;D_4__67-14;Ambiguous_taxa | 1.754557168 | 10 | -1.785868161 | 11 |
| D_0__Bacteria;D_1__Proteobacteria;D_2__Alphaproteobacteria;D_3__Paracaedibacterales;D_4__Paracaedibacteraceae;D_5__uncultured | 0.961024806 | 9 | -0.992335799 | 8 |
| D_0__Bacteria;D_1__Actinobacteria;D_2__Actinobacteria;D_3__Corynebacteriales;D_4__Nocardiaceae;D_5__Gordonia | 0.637663914 | 9 | -0.668974907 | 8 |
| D_0__Bacteria;D_1__Bacteroidetes;D_2__Bacteroidia;D_3__Flavobacteriales;D_4__Crocinitomicaceae;D_5__Salinirepens | 0.301893715 | 9 | -0.333204708 | 8 |
| D_0__Bacteria;D_1__Proteobacteria;D_2__Gammaproteobacteria;D_3__Oceanospirillales;D_4__Saccharospirillaceae;D_5__Oceanobacter | 1.775072851 | 8 | -1.806383844 | 9 |
| D_0__Bacteria;D_1__Proteobacteria;D_2__Gammaproteobacteria;D_3__Pseudomonadales;D_4__Pseudomonadaceae;D_5__Pseudomonas | 1.761579277 | 8 | -1.79289027 | 8 |
| D_0__Bacteria;D_1__Chlamydiae;D_2__Chlamydiae;D_3__Chlamydiales;D_4__Parachlamydiaceae;D_5__Neochlamydia | 1.566372496 | 8 | -1.597683489 | 8 |
| D_0__Bacteria;D_1__Actinobacteria;D_2__Acidimicrobiia;__;__;__ | 0.893835686 | 8 | -0.925146679 | 8 |
| D_0__Bacteria;D_1__Nitrospirae;D_2__Nitrospira;D_3__Nitrospirales;D_4__Nitrospiraceae;D_5__Nitrospira | 0.529545569 | 8 | -0.560856562 | 7 |
| D_0__Bacteria;D_1__Actinobacteria;D_2__Actinobacteria;D_3__Micrococcales;D_4__Dermacoccaceae;D_5__Dermacoccus | 0.373243935 | 8 | -0.404554928 | 7 |

| | | | | |
|---|---|---|---|---|
| D_0__Bacteria;D_1__Proteobacteria;D_2__Deltaproteobacteria;D_3__Myxococcales;D_4__Phaselicystidaceae;D_5__Phaselicystis | 0.25436589 | 8 | -0.285676882 | 7 |
| D_0__Bacteria;D_1__Actinobacteria;D_2__Actinobacteria;D_3__Propionibacteriales;D_4__Nocardioidaceae;D_5__Aeromicrobium | 0.109948532 | 8 | -0.141259525 | 7 |
| D_0__Bacteria;D_1__Proteobacteria;D_2__Alphaproteobacteria;__;__;__ | 0.061747557 | 8 | -0.09305855 | 7 |
| D_0__Bacteria;D_1__Proteobacteria;D_2__Gammaproteobacteria;D_3__Alteromonadales;D_4__Pseudoalteromonadaceae;D_5__Pseudoalteromonas | 0.058334288 | 8 | -0.089645281 | 7 |
| D_0__Bacteria;D_1__Bacteroidetes;D_2__Bacteroidia;D_3__Cytophagales;D_4__Cyclobacteriaceae;D_5__uncultured | 0.029760535 | 8 | -0.061071527 | 7 |
| D_0__Bacteria;D_1__Proteobacteria;D_2__Deltaproteobacteria;D_3__Bdellovibrionales;D_4__Bdellovibrionaceae;D_5__Bdellovibrio | -0.176936911 | 8 | 0.145625918 | 7 |
| D_0__Bacteria;D_1__Verrucomicrobia;D_2__Verrucomicrobiae;D_3__Pedosphaerales;D_4__Pedosphaeraceae;__ | -0.197296649 | 8 | 0.165985657 | 7 |
| D_0__Bacteria;D_1__Proteobacteria;D_2__Alphaproteobacteria;D_3__Rickettsiales;D_4__Anaplasmataceae;D_5__Neorickettsia | -0.213416351 | 8 | 0.182105358 | 7 |
| D_0__Bacteria;D_1__Proteobacteria;D_2__Alphaproteobacteria;D_3__Rhodospirillales;D_4__Rhodospirillaceae;D_5__uncultured | -0.244154859 | 8 | 0.212843866 | 7 |
| D_0__Bacteria;D_1__Bacteroidetes;D_2__Bacteroidia;D_3__Cytophagales;D_4__Spirosomaceae;__ | -0.25084514 | 8 | 0.219534147 | 7 |
| D_0__Bacteria;D_1__Bacteroidetes;D_2__Bacteroidia;D_3__Flavobacteriales;D_4__Flavobacteriaceae;__ | 1.231243224 | 7 | -1.262554217 | 7 |
| D_0__Bacteria;D_1__Proteobacteria;D_2__Gammaproteobacteria;D_3__Betaproteobacteriales;D_4__Burkholderiaceae;D_5__Limnobacter | 0.930876958 | 7 | -0.96218795 | 7 |
| D_0__Bacteria;D_1__Proteobacteria;D_2__Alphaproteobacteria;D_3__Reyranellales;D_4__Reyranellaceae;D_5__Reyranella | 0.573168823 | 7 | -0.604479816 | 6 |
| D_0__Bacteria;D_1__Proteobacteria;D_2__Gammaproteobacteria;D_3__Betaproteobacteriales;D_4__Methylophilaceae;D_5__Methylotenera | 0.50221267 | 7 | -0.533523663 | 6 |
| D_0__Bacteria;D_1__Proteobacteria;D_2__Gammaproteobacteria;D_3__Alteromonadales;D_4__Marinobacteraceae;D_5__Marinobacter | 0.455484585 | 7 | -0.486795578 | 6 |
| D_0__Bacteria;D_1__Proteobacteria;D_2__Gammaproteobacteria;D_3__Oceanospirillales;D_4__Pseudohongiellaceae;D_5__Pseudohongiella | 0.378437878 | 7 | -0.409748871 | 6 |
| D_0__Bacteria;D_1__Proteobacteria;D_2__Gammaproteobacteria;D_3__Oceanospirillales;D_4__Alcanivoracaceae;D_5__Alcanivorax | 0.222942653 | 7 | -0.254253646 | 6 |
| D_0__Bacteria;D_1__Bacteroidetes;D_2__Bacteroidia;D_3__Sphingobacteriales;D_4__NS11-12 marine group;D_5__uncultured bacterium | 0.156442248 | 7 | -0.187753241 | 6 |
| D_0__Bacteria;D_1__Proteobacteria;D_2__Gammaproteobacteria;D_3__Salinisphaerales;D_4__Solimonadaceae;D_5__Nevskia | 0.120955263 | 7 | -0.152266256 | 6 |
| D_0__Bacteria;D_1__Actinobacteria;D_2__Actinobacteria;D_3__Micrococcales;D_4__Brevibacteriaceae;D_5__Brevibacterium | 0.094353202 | 7 | -0.125664195 | 6 |
| D_0__Bacteria;D_1__Planctomycetes;D_2__Planctomycetacia;D_3__Gemmatales;D_4__Gemmataceae;D_5__Fimbriiglobus | 0.042042956 | 7 | -0.073353949 | 6 |
| D_0__Bacteria;D_1__Proteobacteria;D_2__Gammaproteobacteria;D_3__Betaproteobacteriales;D_4__Nitrosomonadaceae;D_5__Nitrosomonas | 0.034336976 | 7 | -0.065647969 | 6 |
| D_0__Bacteria;D_1__Planctomycetes;D_2__OM190;__;__;__ | -0.028365716 | 7 | -0.002945277 | 6 |
| D_0__Bacteria;D_1__Actinobacteria;D_2__Actinobacteria;D_3__Micrococcales;D_4__Micrococcaceae;D_5__Rothia | -0.036487415 | 7 | 0.005176422 | 6 |
| D_0__Bacteria;D_1__Proteobacteria;D_2__Gammaproteobacteria;D_3__Pseudomonadales;D_4__Moraxellaceae;D_5__[Agitococcus] lubricus group | -0.067623414 | 7 | 0.036312422 | 6 |
| D_0__Bacteria;D_1__Actinobacteria;D_2__Actinobacteria;D_3__Corynebacteriales;D_4__Nocardiaceae;D_5__Williamsia | -0.072484785 | 7 | 0.041173792 | 6 |
| D_0__Bacteria;D_1__Gemmatimonadetes;D_2__Gemmatimonadetes;D_3__Gemmatimonadales;D_4__Gemmatimonadaceae;D_5__uncultured | -0.078789833 | 7 | 0.04747884 | 6 |
| D_0__Bacteria;D_1__Planctomycetes;D_2__Planctomycetacia;D_3__Planctomycetales;__;__ | -0.079601165 | 7 | 0.048290172 | 6 |
| D_0__Bacteria;D_1__Proteobacteria;D_2__Gammaproteobacteria;D_3__Ga0077536;Ambiguous_taxa;Ambiguous_taxa | -0.096338402 | 7 | 0.065027409 | 6 |
| D_0__Bacteria;D_1__Proteobacteria;D_2__Alphaproteobacteria;D_3__Rhizobiales;D_4__Xanthobacteraceae;D_5__Pseudorhodoplanes | -0.103204301 | 7 | 0.071893308 | 6 |
| D_0__Bacteria;D_1__Planctomycetes;D_2__Planctomycetacia;D_3__Isosphaerales;D_4__Isosphaeraceae;D_5__Singulisphaera | -0.121887525 | 7 | 0.090576532 | 6 |
| D_0__Bacteria;D_1__Actinobacteria;D_2__Actinobacteria;D_3__Micrococcales;D_4__Dermatophilaceae;D_5__Mobilicoccus | -0.124889317 | 7 | 0.093578324 | 6 |
| D_0__Bacteria;D_1__Firmicutes;D_2__Clostridia;D_3__Thermoanaerobacterales;D_4__Family III;D_5__Thermoanaerobacterium | -0.127628457 | 7 | 0.096317464 | 6 |

| | | | | |
|---|---|---|---|---|
| D_0__Bacteria;D_1__Bacteroidetes;D_2__Bacteroidia;D_3__Cytophagales;D_4__Microscillaceae;D_5__uncultured | -0.127823067 | 7 | 0.096512075 | 6 |
| D_0__Bacteria;D_1__Hydrogenedentes;D_2__Hydrogenedentia;D_3__Hydrogenedentiales;D_4__Hydrogenedensaceae;D_5__metagenome | -0.138933876 | 7 | 0.107622883 | 6 |
| D_0__Bacteria;D_1__Verrucomicrobia;D_2__Verrucomicrobiae;D_3__Verrucomicrobiales;D_4__DEV007;__ | -0.141112608 | 7 | 0.109801615 | 6 |
| D_0__Bacteria;D_1__Bacteroidetes;D_2__Bacteroidia;D_3__Chitinophagales;D_4__Saprospiraceae;__ | -0.144697508 | 7 | 0.113386515 | 6 |
| D_0__Bacteria;D_1__Planctomycetes;D_2__OM190;D_3__uncultured bacterium;D_4__uncultured bacterium;D_5__uncultured bacterium | -0.149132154 | 7 | 0.117821161 | 6 |
| D_0__Bacteria;D_1__Bacteroidetes;D_2__Bacteroidia;D_3__Flavobacteriales;D_4__Cryomorphaceae;D_5__Owenweeksia | -0.150542029 | 7 | 0.119231036 | 6 |
| D_0__Bacteria;D_1__Proteobacteria;D_2__Alphaproteobacteria;D_3__Rhodospirillales;D_4__Terasakiellaceae;D_5__uncultured | -0.152717054 | 7 | 0.121406061 | 6 |
| D_0__Bacteria;D_1__Proteobacteria;D_2__Gammaproteobacteria;D_3__Gammaproteobacteria Incertae Sedis;D_4__Unknown Family;Ambiguous_taxa | -0.155627825 | 7 | 0.124316832 | 6 |
| D_0__Bacteria;D_1__Bacteroidetes;D_2__Bacteroidia;D_3__Cytophagales;D_4__Cytophagaceae;D_5__Cytophaga | -0.158078069 | 7 | 0.126767076 | 6 |
| D_0__Bacteria;D_1__Chloroflexi;D_2__Chloroflexia;D_3__Thermomicrobiales;D_4__JG30-KF-CM45;D_5__Paraburkholderia tropica | -0.159407335 | 7 | 0.128096342 | 6 |
| D_0__Bacteria;D_1__Firmicutes;D_2__Bacilli;D_3__Lactobacillales;D_4__Carnobacteriaceae;D_5__Carnobacterium | -0.159407335 | 7 | 0.128096342 | 6 |
| D_0__Bacteria;D_1__Proteobacteria;D_2__Alphaproteobacteria;D_3__Rhizobiales;D_4__Beijerinckiaceae;__ | -0.167080766 | 7 | 0.135769773 | 6 |
| D_0__Bacteria;D_1__Proteobacteria;D_2__Alphaproteobacteria;D_3__Holosporales;D_4__Holosporaceae;D_5__uncultured | -0.168162627 | 7 | 0.136851634 | 6 |
| D_0__Bacteria;D_1__Bacteroidetes;D_2__Bacteroidia;D_3__Flavobacteriales;D_4__Flavobacteriaceae;D_5__Cellulophaga | -0.174486667 | 7 | 0.143175674 | 6 |
| D_0__Bacteria;D_1__Bacteroidetes;D_2__Bacteroidia;D_3__Flavobacteriales;D_4__Weeksellaceae;D_5__Chryseobacterium | -0.176936911 | 7 | 0.145625918 | 6 |
| D_0__Bacteria;D_1__Proteobacteria;D_2__Alphaproteobacteria;D_3__Rickettsiales;D_4__Midichloriaceae;D_5__Lyticum | -0.177592048 | 7 | 0.146281055 | 6 |
| D_0__Bacteria;D_1__Proteobacteria;D_2__Gammaproteobacteria;D_3__Alteromonadales;D_4__Idiomarinaceae;D_5__Idiomarina | -0.179676052 | 7 | 0.148365059 | 6 |
| D_0__Bacteria;D_1__Dependentiae;D_2__Babeliae;D_3__Babeliales;D_4__UBA12409;D_5__metagenome | -0.180042293 | 7 | 0.1487313 | 6 |
| D_0__Bacteria;D_1__Actinobacteria;D_2__Actinobacteria;D_3__Micrococcales;D_4__Promicromonosporaceae;D_5__Cellulosimicrobium | -0.182781433 | 7 | 0.15147044 | 6 |
| D_0__Bacteria;D_1__Bacteroidetes;D_2__Bacteroidia;D_3__Flavobacteriales;D_4__Flavobacteriaceae;D_5__Mesonia | -0.191799236 | 7 | 0.160488243 | 6 |
| D_0__Bacteria;D_1__Proteobacteria;D_2__Alphaproteobacteria;D_3__Rhodospirillales;D_4__uncultured;D_5__uncultured soil bacterium | -0.198085058 | 7 | 0.166774065 | 6 |
| D_0__Bacteria;D_1__Proteobacteria;D_2__Gammaproteobacteria;D_3__Pasteurellales;D_4__Pasteurellaceae;D_5__Haemophilus | -0.204509555 | 7 | 0.173198562 | 6 |
| D_0__Bacteria;D_1__Proteobacteria;D_2__Alphaproteobacteria;D_3__Rhizobiales;D_4__Beijerinckiaceae;D_5__Bosea | -0.205591416 | 7 | 0.174280423 | 6 |
| D_0__Bacteria;D_1__Planctomycetes;D_2__Planctomycetacia;D_3__Pirellulales;D_4__Pirellulaceae;D_5__Blastopirellula | -0.215020837 | 7 | 0.183709844 | 6 |
| D_0__Bacteria;D_1__Proteobacteria;D_2__Alphaproteobacteria;D_3__Caulobacterales;D_4__Hyphomonadaceae;D_5__Hyphomonas | -0.238965474 | 7 | 0.207654481 | 6 |
| D_0__Bacteria;D_1__Firmicutes;D_2__Clostridia;D_3__Clostridiales;D_4__Clostridiaceae 1;D_5__Clostridium sensu stricto 1 | -1.589713093 | 7 | 1.558402101 | 6 |
| D_0__Bacteria;D_1__Proteobacteria;D_2__Gammaproteobacteria;D_3__Betaproteobacteriales;D_4__Burkholderiaceae;D_5__Undibacterium | 1.037452226 | 6 | -1.068763219 | 5 |
| D_0__Bacteria;D_1__Actinobacteria;D_2__Acidimicrobiia;D_3__Microtrichales;D_4__Iamiaceae;D_5__Iamia | 0.954663671 | 6 | -0.985974664 | 6 |
| D_0__Bacteria;D_1__Proteobacteria;D_2__Alphaproteobacteria;D_3__Micavibrionales;D_4__Micavibrionaceae;D_5__uncultured | 0.93887395 | 6 | -0.970184943 | 5 |
| D_0__Bacteria;D_1__Planctomycetes;D_2__Planctomycetacia;D_3__Planctomycetales;D_4__Rubinisphaeraceae;D_5__uncultured | 0.909667526 | 6 | -0.940978519 | 5 |
| D_0__Bacteria;D_1__Actinobacteria;D_2__Thermoleophilia;D_3__Solirubrobacterales;D_4__Solirubrobacteraceae;D_5__Parviterribacter | 0.727234 | 6 | -0.758544993 | 6 |
| D_0__Bacteria;D_1__Actinobacteria;D_2__Actinobacteria;D_3__Corynebacteriales;D_4__Nocardiaceae;D_5__Nocardia | 0.59614175 | 6 | -0.627452743 | 5 |
| D_0__Bacteria;D_1__Bacteroidetes;D_2__Bacteroidia;D_3__Cytophagales;D_4__Microscillaceae;D_5__OLB12 | 0.494641884 | 6 | -0.525952876 | 5 |

| | | | | |
|---|---|---|---|---|
| D_0__Bacteria;D_1__Firmicutes;D_2__Bacilli;D_3__Lactobacillales;D_4__Enterococcaceae;D_5__Enterococcus | 0.39432836 | 6 | -0.425639353 | 5 |
| D_0__Bacteria;D_1__Bacteroidetes;D_2__Bacteroidia;D_3__Chitinophagales;D_4__Saprospiraceae;D_5__Aureispira | 0.33717656 | 6 | -0.368487553 | 5 |
| D_0__Bacteria;D_1__Proteobacteria;D_2__Alphaproteobacteria;D_3__Sphingomonadales;D_4__Sphingomonadaceae;D_5__Sphingopyxis | 0.155339516 | 6 | -0.186650509 | 5 |
| D_0__Bacteria;D_1__Proteobacteria;D_2__Gammaproteobacteria;D_3__Oceanospirillales;D_4__Halomonadaceae;D_5__Salinicola | 0.138888008 | 6 | -0.170199001 | 5 |
| D_0__Bacteria;D_1__Actinobacteria;D_2__Actinobacteria;D_3__Micrococcales;D_4__Intrasporangiaceae;__ | 0.124696515 | 6 | -0.156007507 | 5 |
| D_0__Bacteria;D_1__Proteobacteria;D_2__Gammaproteobacteria;D_3__Cellvibrionales;D_4__Cellvibrionaceae;D_5__Cellvibrio | 0.112351557 | 6 | -0.14366255 | 5 |
| D_0__Bacteria;D_1__Proteobacteria;D_2__Alphaproteobacteria;D_3__Rhizobiales;D_4__Rhizobiales Incertae Sedis;D_5__Bauldia | 0.082971301 | 6 | -0.114282294 | 5 |
| D_0__Bacteria;D_1__Proteobacteria;D_2__Gammaproteobacteria;D_3__Salinisphaerales;D_4__Solimonadaceae;D_5__Polycyclovorans | 0.061394684 | 6 | -0.092705677 | 5 |
| D_0__Bacteria;D_1__Proteobacteria;D_2__Gammaproteobacteria;D_3__Betaproteobacteriales;D_4__Rhodocyclaceae;D_5__Methyloversatilis | -0.038793394 | 6 | 0.007482401 | 5 |
| D_0__Bacteria;D_1__Proteobacteria;D_2__Gammaproteobacteria;D_3__Betaproteobacteriales;D_4__Burkholderiaceae;D_5__Simplicispira | -0.062014995 | 6 | 0.030704002 | 5 |
| D_0__Bacteria;D_1__Proteobacteria;D_2__Gammaproteobacteria;D_3__Salinisphaerales;D_4__Solimonadaceae;D_5__uncultured | -0.088713312 | 6 | 0.057402319 | 5 |
| D_0__Bacteria;D_1__Proteobacteria;D_2__Alphaproteobacteria;D_3__Sphingomonadales;D_4__Sphingomonadaceae;D_5__Sphingomonas | -0.100773049 | 6 | 0.069462056 | 5 |
| D_0__Bacteria;D_1__Proteobacteria;D_2__Gammaproteobacteria;D_3__Betaproteobacteriales;D_4__Burkholderiaceae;D_5__Ralstonia | -0.104529579 | 6 | 0.073218586 | 5 |
| D_0__Bacteria;D_1__Bacteroidetes;D_2__Bacteroidia;D_3__Flavobacteriales;D_4__Flavobacteriaceae;D_5__Zunongwangia | -0.124889317 | 6 | 0.093578324 | 5 |
| D_0__Bacteria;D_1__Proteobacteria;D_2__Gammaproteobacteria;D_3__Betaproteobacteriales;D_4__Rhodocyclaceae;D_5__Zoogloea | -0.173630528 | 6 | 0.142319535 | 5 |
| D_0__Bacteria;D_1__Proteobacteria;D_2__Deltaproteobacteria;D_3__Bdellovibrionales;D_4__Bacteriovoracaceae;D_5__Peredibacter | -0.483236603 | 6 | 0.45192561 | 5 |
| D_0__Bacteria;D_1__Proteobacteria;D_2__Alphaproteobacteria;D_3__Caulobacterales;D_4__Hyphomonadaceae;D_5__SWB02 | -0.490691077 | 6 | 0.459380084 | 5 |
| D_0__Bacteria;D_1__Proteobacteria;D_2__Alphaproteobacteria;D_3__Caulobacterales;D_4__Caulobacteraceae;D_5__Caulobacter | -0.532359901 | 6 | 0.501048909 | 5 |
| D_0__Bacteria;D_1__Actinobacteria;D_2__Actinobacteria;D_3__Micrococcales;D_4__Micrococcaceae;__ | -0.595613926 | 6 | 0.564302933 | 5 |
| D_0__Bacteria;D_1__Proteobacteria;D_2__Gammaproteobacteria;D_3__Betaproteobacteriales;D_4__Burkholderiaceae;D_5__Cupriavidus | -0.60533499 | 6 | 0.574023998 | 5 |
| D_0__Bacteria;D_1__Spirochaetes;D_2__Leptospirae;D_3__Leptospirales;D_4__Leptospiraceae;D_5__Turneriella | -0.606823516 | 6 | 0.575512524 | 5 |
| D_0__Bacteria;D_1__Actinobacteria;D_2__Actinobacteria;D_3__Actinomycetales;D_4__Actinomycetaceae;D_5__Actinomyces | -0.608148728 | 6 | 0.576837736 | 5 |
| D_0__Bacteria;D_1__Deinococcus-Thermus;D_2__Deinococci;D_3__Deinococcales;D_4__Deinococcaceae;D_5__Deinococcus | -0.680542968 | 6 | 0.649231975 | 5 |
| D_0__Bacteria;D_1__Actinobacteria;D_2__Actinobacteria;D_3__Corynebacteriales;D_4__Nocardiaceae;D_5__Millisia | 1.452683679 | 5 | -1.483994672 | 5 |
| D_0__Bacteria;D_1__Actinobacteria;D_2__Acidimicrobiia;D_3__Microtrichales;D_4__Microtrichaceae;D_5__IMCC26207 | 1.027146054 | 5 | -1.058457047 | 5 |
| D_0__Bacteria;D_1__Actinobacteria;D_2__Actinobacteria;D_3__Micrococcales;D_4__Micrococcaceae;D_5__Kocuria | 0.847889638 | 5 | -0.879200631 | 5 |
| D_0__Bacteria;D_1__Bacteroidetes;D_2__Bacteroidia;D_3__Cytophagales;D_4__Cyclobacteriaceae;D_5__Roseivirga | 0.352156625 | 5 | NA | NA |
| D_0__Bacteria;D_1__Proteobacteria;D_2__Gammaproteobacteria;__;__;__ | 0.231597152 | 5 | -0.262908145 | 4 |
| D_0__Bacteria;D_1__Proteobacteria;D_2__Alphaproteobacteria;D_3__Rhizobiales;D_4__Rhizobiaceae;__ | 0.116348074 | 5 | -0.147659067 | 4 |
| D_0__Bacteria;D_1__Proteobacteria;D_2__Alphaproteobacteria;D_3__Rickettsiales;D_4__Rickettsiaceae;D_5__uncultured | 0.069958642 | 5 | -0.101269635 | 4 |
| D_0__Bacteria;D_1__Actinobacteria;D_2__Actinobacteria;D_3__Frankiales;D_4__Nakamurellaceae;D_5__Nakamurella | -0.245767466 | 5 | 0.214456473 | 4 |
| D_0__Bacteria;D_1__Bacteroidetes;D_2__Bacteroidia;D_3__Cytophagales;D_4__Spirosomaceae;D_5__Taeseokella | -0.246530195 | 5 | 0.215219203 | 4 |
| D_0__Bacteria;D_1__Bacteroidetes;D_2__Bacteroidia;D_3__Flavobacteriales;D_4__Flavobacteriaceae;D_5__Muricauda | -0.890426769 | 5 | 0.859115776 | 4 |

| | | | | |
|---|---|---|---|---|
| D_0__Bacteria;D_1__Proteobacteria;D_2__Alphaproteobacteria;D_3__Rhizobiales;D_4__Hyphomicrobiaceae;D_5__Filomicrobium | -0.909702367 | 5 | 0.878391374 | 4 |
| D_0__Bacteria;D_1__Proteobacteria;D_2__Alphaproteobacteria;D_3__Rhizobiales;D_4__Rhizobiaceae;D_5__Shinella | -1.223220811 | 5 | 1.191909818 | 4 |
| D_0__Bacteria;D_1__Chlamydiae;D_2__Chlamydiae;D_3__Chlamydiales;D_4__Parachlamydiaceae;D_5__uncultured | 0.670680608 | 4 | -0.701991601 | 4 |
| D_0__Bacteria;D_1__Firmicutes;D_2__Bacilli;D_3__Lactobacillales;D_4__Streptococcaceae;D_5__Streptococcus | 0.502410389 | 4 | -0.533721382 | 4 |
| D_0__Bacteria;D_1__Proteobacteria;D_2__Gammaproteobacteria;D_3__Pseudomonadales;D_4__Moraxellaceae;D_5__Psychrobacter | 0.194154875 | 4 | -0.225465868 | 3 |
| D_0__Bacteria;D_1__Actinobacteria;D_2__Actinobacteria;D_3__Micrococcales;D_4__Microbacteriaceae;D_5__Microbacterium | 0.131018802 | 4 | -0.162329795 | 4 |
| D_0__Bacteria;D_1__Planctomycetes;D_2__Planctomycetacia;D_3__Pirellulales;D_4__Pirellulaceae;D_5__Pirellula | 0.053577665 | 4 | -0.084888658 | 3 |
| D_0__Bacteria;D_1__Bacteroidetes;D_2__Bacteroidia;D_3__Flavobacteriales;D_4__Crocinitomicaceae;D_5__Fluviicola | -0.014556762 | 4 | -0.016754231 | 3 |
| D_0__Bacteria;D_1__Proteobacteria;D_2__Alphaproteobacteria;D_3__Caulobacterales;D_4__Hyphomonadaceae;D_5__Hirschia | -0.054712697 | 4 | 0.023401704 | 3 |
| D_0__Bacteria;D_1__Proteobacteria;D_2__Alphaproteobacteria;D_3__Rhodobacterales;D_4__Rhodobacteraceae;D_5__Amaricoccus | -0.133994786 | 4 | 0.102683793 | 3 |
| D_0__Bacteria;D_1__Chloroflexi;D_2__Anaerolineae;D_3__Caldilineales;D_4__Caldilineaceae;D_5__uncultured | -0.288578258 | 4 | 0.257267265 | 3 |
| D_0__Bacteria;D_1__Proteobacteria;D_2__Gammaproteobacteria;D_3__Vibrionales;D_4__Vibrionaceae;D_5__Vibrio | -0.524040765 | 4 | 1.800451397 | 7 |
| D_0__Bacteria;D_1__Proteobacteria;D_2__Gammaproteobacteria;D_3__Cellvibrionales;D_4__Halieaceae;D_5__Marimicrobium | -0.592119523 | 4 | 0.56080853 | 3 |
| D_0__Bacteria;D_1__Proteobacteria;D_2__Alphaproteobacteria;D_3__Caulobacterales;D_4__Caulobacteraceae;D_5__Brevundimonas | -0.658141414 | 4 | 0.626830421 | 3 |
| D_0__Bacteria;D_1__Proteobacteria;D_2__Alphaproteobacteria;D_3__Rhizobiales;D_4__Rhizobiaceae;D_5__Allorhizobium-Neorhizobium-Pararhizobium-Rhizobium | -1.412179024 | 4 | 1.380868031 | 3 |
| D_0__Bacteria;D_1__Proteobacteria;D_2__Gammaproteobacteria;D_3__Alteromonadales;D_4__Alteromonadaceae;__ | 1.373507101 | 3 | -1.404818094 | 3 |
| D_0__Bacteria;D_1__Proteobacteria;D_2__Alphaproteobacteria;D_3__Rhodobacterales;D_4__Rhodobacteraceae;D_5__Pseudorhodobacter | 1.063645794 | 3 | -1.094956787 | 3 |
| D_0__Bacteria;D_1__Proteobacteria;D_2__Alphaproteobacteria;D_3__Rhizobiales;D_4__Xanthobacteraceae;__ | 0.215866828 | 3 | -0.247177821 | 2 |
| D_0__Bacteria;D_1__Firmicutes;D_2__Bacilli;D_3__Bacillales;D_4__Bacillaceae;D_5__Bacillus | -0.631453825 | 3 | 0.600142832 | 2 |
| D_0__Bacteria;D_1__Actinobacteria;D_2__Actinobacteria;D_3__Corynebacteriales;D_4__Corynebacteriaceae;D_5__Corynebacterium 1 | -0.779596427 | 3 | 0.748285434 | 2 |
| D_0__Bacteria;D_1__Planctomycetes;D_2__Planctomycetacia;D_3__Pirellulales;D_4__Pirellulaceae;D_5__Rhodopirellula | -0.859914041 | 3 | 0.828603048 | 2 |
| D_0__Bacteria;D_1__Proteobacteria;D_2__Alphaproteobacteria;D_3__Rhizobiales;D_4__Stappiaceae;D_5__Stappia | -1.052777031 | 3 | 1.021466038 | 2 |
| D_0__Bacteria;D_1__Proteobacteria;D_2__Gammaproteobacteria;D_3__Betaproteobacteriales;D_4__Burkholderiaceae;D_5__Hydrogenophaga | 0.687175701 | 2 | -0.718486694 | 1 |
| D_0__Bacteria;D_1__Bacteroidetes;D_2__Bacteroidia;D_3__Flavobacteriales;D_4__Flavobacteriaceae;D_5__Flavobacterium | 0.656188819 | 2 | -0.687499812 | 1 |
| D_0__Bacteria;D_1__Actinobacteria;D_2__Actinobacteria;D_3__Micrococcales;D_4__Microbacteriaceae;D_5__Pseudoclavibacter | 0.601321561 | 2 | -0.632632554 | 2 |
| D_0__Bacteria;D_1__Proteobacteria;D_2__Gammaproteobacteria;D_3__Enterobacteriales;D_4__Enterobacteriaceae;__ | 0.302018433 | 2 | -0.333329426 | 2 |
| D_0__Bacteria;D_1__Proteobacteria;D_2__Gammaproteobacteria;D_3__Betaproteobacteriales;D_4__Burkholderiaceae;__ | 0.172202309 | 2 | -0.203513302 | 1 |
| D_0__Bacteria;D_1__Chloroflexi;D_2__Chloroflexia;D_3__Thermomicrobiales;D_4__JG30-KF-CM45;Ambiguous_taxa | 0.118414119 | 2 | -0.149725112 | 1 |
| D_0__Bacteria;D_1__Firmicutes;D_2__Bacilli;D_3__Bacillales;D_4__Staphylococcaceae;D_5__Staphylococcus | -0.173361148 | 2 | 0.142050155 | 1 |
| D_0__Bacteria;D_1__Proteobacteria;D_2__Alphaproteobacteria;D_3__Rhodobacterales;D_4__Rhodobacteraceae;D_5__Paracoccus | -0.237129954 | 2 | 0.205818962 | 1 |
| D_0__Bacteria;D_1__Bacteroidetes;D_2__Bacteroidia;D_3__Flavobacteriales;D_4__Flavobacteriaceae;D_5__Arenibacter | -0.257267489 | 2 | 0.225956496 | 1 |
| D_0__Bacteria;D_1__Proteobacteria;D_2__Alphaproteobacteria;D_3__Rhizobiales;D_4__Xanthobacteraceae;D_5__Pseudoxanthobacter | -0.384489415 | 2 | 0.353178422 | 1 |
| D_0__Bacteria;D_1__Proteobacteria;D_2__Alphaproteobacteria;D_3__Sphingomonadales;D_4__Sphingomonadaceae;D_5__Sphingobium | -0.399005926 | 2 | 0.367694933 | 1 |

| | | | | |
|---|---|---|---|---|
| D_0__Bacteria;D_1__Planctomycetes;D_2__Planctomycetacia;D_3__Pirellulales;D_4__Pirellulaceae;D_5__Pir4 lineage | -0.478195495 | 2 | 0.446884502 | 1 |
| D_0__Bacteria;D_1__Proteobacteria;D_2__Alphaproteobacteria;D_3__Rhodobacterales;D_4__Rhodobacteraceae;D_5__Defluviimonas | -0.527466452 | 2 | 0.496155459 | 1 |
| D_0__Bacteria;D_1__Proteobacteria;D_2__Alphaproteobacteria;D_3__Rhizobiales;__;__ | -0.591566827 | 2 | 0.560255834 | 1 |
| D_0__Bacteria;D_1__Actinobacteria;D_2__Actinobacteria;D_3__Pseudonocardiales;D_4__Pseudonocardiaceae;D_5__Pseudonocardia | -0.705904992 | 2 | 0.674593999 | 1 |
| D_0__Bacteria;D_1__Bacteroidetes;D_2__Bacteroidia;D_3__Flavobacteriales;D_4__Flavobacteriaceae;D_5__Maribacter | -0.713065014 | 2 | 0.681754021 | 1 |
| D_0__Bacteria;D_1__Proteobacteria;D_2__Alphaproteobacteria;D_3__Rhizobiales;D_4__Devosiaceae;D_5__uncultured | -0.789179017 | 2 | 0.757868024 | 1 |
| D_0__Bacteria;D_1__Planctomycetes;D_2__Planctomycetacia;D_3__Planctomycetales;D_4__Gimesiaceae;D_5__uncultured | -0.829450488 | 2 | 0.798139495 | 1 |
| D_0__Bacteria;D_1__Proteobacteria;D_2__Alphaproteobacteria;D_3__Rhizobiales;D_4__Devosiaceae;D_5__Devosia | -1.437212661 | 2 | 1.405901669 | 1 |
| D_0__Bacteria;D_1__Actinobacteria;D_2__Actinobacteria;D_3__Corynebacteriales;D_4__Nocardiaceae;D_5__Rhodococcus | 0.514157267 | 1 | -0.54546826 | 1 |
| D_0__Bacteria;D_1__Actinobacteria;D_2__Acidimicrobiia;D_3__Microtrichales;D_4__uncultured;D_5__uncultured bacterium | 0.466513188 | 1 | -0.497824181 | 1 |
| D_0__Bacteria;D_1__Proteobacteria;D_2__Gammaproteobacteria;D_3__Betaproteobacteriales;D_4__Chitinibacteraceae;D_5__Deefgea | 0.369365492 | 1 | -0.400676485 | 0 |
| D_0__Bacteria;D_1__Proteobacteria;D_2__Deltaproteobacteria;D_3__Myxococcales;D_4__Nannocystaceae;D_5__Nannocystis | 0.15840122 | 1 | -0.189712213 | 0 |
| D_0__Bacteria;D_1__Proteobacteria;D_2__Alphaproteobacteria;D_3__Rhizobiales;D_4__Hyphomicrobiaceae;D_5__Hyphomicrobium | 0.106433126 | 1 | -0.137744119 | 1 |
| D_0__Bacteria;D_1__Proteobacteria;D_2__Alphaproteobacteria;D_3__Rhizobiales;D_4__Hyphomicrobiaceae;D_5__Pedomicrobium | -0.145620615 | 1 | 0.114309622 | 0 |
| D_0__Bacteria;D_1__Proteobacteria;D_2__Gammaproteobacteria;D_3__Legionellales;D_4__Legionellaceae;D_5__Legionella | -0.311806618 | 1 | 0.280495626 | 0 |
| D_0__Bacteria;D_1__Proteobacteria;D_2__Alphaproteobacteria;D_3__Rhizobiales;D_4__Rhizobiaceae;D_5__Mesorhizobium | -0.525241475 | 1 | 0.493930482 | 0 |
| D_0__Bacteria;D_1__Proteobacteria;D_2__Alphaproteobacteria;D_3__Rhizobiales;D_4__Rhizobiaceae;D_5__Hoeflea | -0.804972616 | 1 | 0.773661623 | 0 |
| D_0__Bacteria;D_1__Proteobacteria;D_2__Gammaproteobacteria;D_3__Enterobacteriales;D_4__Enterobacteriaceae;D_5__Hafnia-Obesumbacterium | 0.371621782 | 0 | -0.402932775 | 0 |
| D_0__Bacteria;D_1__Bacteroidetes;D_2__Bacteroidia;D_3__Cytophagales;D_4__Amoebophilaceae;D_5__Candidatus Amoebophilus | 0.060877796 | 0 | -0.092188789 | 0 |

**Table S3:** Test statistics of differential abundance analysis between fecal samples before and after the diet change in the second experiment using ANCOM in QIIME2 on the phylum level, with and without filtering of diet-derived bacteria. Test statistics of bacterial phyla that significantly differ between diet treatments are indicated in bold.

| | no diet filtering | | diet filtering | |
|---|---|---|---|---|
| pyhlum | clr | W | clr | W |
| D_0__Bacteria;D_1__Firmicutes | **3.95493427** | **11** | **-2.96915209** | **9** |
| D_0__Bacteria;D_1__Chloroflexi | **3.04119235** | **10** | **-3.13592011** | **9** |
| D_0__Bacteria;D_1__RsaHF231 | **2.53840877** | **10** | **-2.63313653** | **9** |
| D_0__Bacteria;D_1__Planctomycetes | 0.21430583 | 8 | **-0.30903359** | **8** |
| D_0__Bacteria;D_1__Actinobacteria | -1.60437486 | 5 | 1.5096471 | 5 |
| D_0__Bacteria;D_1__Proteobacteria | -2.18964059 | 4 | 2.16332499 | 5 |
| D_0__Bacteria;D_1__Deinococcus-Thermus | -1.5918736 | 4 | 1.49714584 | 4 |
| D_0__Bacteria;D_1__Verrucomicrobia | -1.4964669 | 4 | 1.40173914 | 4 |
| D_0__Bacteria;D_1__Fusobacteria | -1.45498136 | 4 | 1.3602536 | 4 |
| D_0__Bacteria;D_1__Bacteroidetes | -0.09079868 | 4 | -0.01611806 | 5 |
| D_0__Bacteria;D_1__Chlamydiae | -0.66459372 | 3 | 0.56986596 | 3 |
| D_0__Bacteria;D_1__Patescibacteria | -0.65611151 | 3 | 0.56138375 | 3 |

**Table S4:** Test statistics of differential abundance analysis between fecal samples before and after the diet change in the second experiment using ANCOM in QIIME2 on the genus level, with and without filtering of diet-derived bacteria. Test statistics of bacterial genera that significantly differ between diet treatments are indicated in bold.

| genus |
|---|
| D_0__Bacteria;D_1__Firmicutes;D_2__Bacilli;D_3__Lactobacillales;D_4__Carnobacteriaceae;D_5__Carnobacterium |
| D_0__Bacteria;D_1__Firmicutes;D_2__Erysipelotrichia;D_3__Erysipelotrichales;D_4__Erysipelotrichaceae;D_5__ZOR0006 |
| D_0__Bacteria;D_1__Firmicutes;D_2__Erysipelotrichia;D_3__Erysipelotrichales;D_4__Erysipelotrichaceae;__ |
| D_0__Bacteria;D_1__Firmicutes;D_2__Clostridia;D_3__Clostridiales;D_4__Peptostreptococcaceae;D_5__Romboutsia |
| D_0__Bacteria;D_1__Proteobacteria;D_2__Gammaproteobacteria;D_3__Vibrionales;D_4__Vibrionaceae;D_5__Vibrio |
| D_0__Bacteria;D_1__Actinobacteria;D_2__Actinobacteria;D_3__Micrococcales;D_4__Microbacteriaceae;D_5__Pseudoclavibacter |
| D_0__Bacteria;D_1__Proteobacteria;D_2__Gammaproteobacteria;D_3__Alteromonadales;D_4__Shewanellaceae;D_5__Shewanella |
| D_0__Bacteria;D_1__Firmicutes;D_2__Clostridia;D_3__Clostridiales;D_4__Eubacteriaceae;D_5__uncultured |
| D_0__Bacteria;D_1__RsaHF231;D_2__uncultured bacterium;D_3__uncultured bacterium;D_4__uncultured bacterium;D_5__uncultured bacterium |
| D_0__Bacteria;D_1__Firmicutes;D_2__Clostridia;D_3__Clostridiales;D_4__Lachnospiraceae;__ |
| D_0__Bacteria;D_1__Firmicutes;D_2__Clostridia;D_3__Clostridiales;D_4__Peptostreptococcaceae;D_5__Intestinibacter |
| D_0__Bacteria;D_1__Firmicutes;D_2__Clostridia;D_3__Clostridiales;D_4__Clostridiaceae 1;D_5__Clostridium sensu stricto 1 |
| D_0__Bacteria;D_1__Firmicutes;D_2__Erysipelotrichia;D_3__Erysipelotrichales;D_4__Erysipelotrichaceae;D_5__Turicibacter |
| D_0__Bacteria;D_1__Chloroflexi;D_2__Chloroflexia;D_3__Thermomicrobiales;D_4__JG30-KF-CM45;D_5__uncultured soil bacterium |
| D_0__Bacteria;D_1__Firmicutes;D_2__Clostridia;D_3__Clostridiales;D_4__Ruminococcaceae;__ |
| D_0__Bacteria;D_1__Actinobacteria;D_2__Actinobacteria;D_3__Actinomycetales;D_4__Actinomycetaceae;D_5__uncultured |
| D_0__Bacteria;D_1__Actinobacteria;D_2__Actinobacteria;D_3__Micrococcales;D_4__Microbacteriaceae;D_5__Leucobacter |
| D_0__Bacteria;D_1__Firmicutes;D_2__Clostridia;D_3__Clostridiales;D_4__Lachnospiraceae;D_5__Tyzzerella 3 |
| D_0__Bacteria;D_1__Chloroflexi;D_2__Chloroflexia;D_3__Thermomicrobiales;D_4__JG30-KF-CM45;D_5__uncultured anaerobic ammonium-oxidizing bacterium |
| D_0__Bacteria;D_1__Actinobacteria;D_2__Actinobacteria;D_3__Bifidobacteriales;D_4__Bifidobacteriaceae;D_5__Bifidobacterium |
| D_0__Bacteria;D_1__Firmicutes;D_2__Clostridia;D_3__Clostridiales;D_4__Peptostreptococcaceae;D_5__Terrisporobacter |
| D_0__Bacteria;D_1__Actinobacteria;D_2__Thermoleophilia;D_3__Solirubrobacterales;D_4__Solirubrobacteraceae;D_5__Conexibacter |
| D_0__Bacteria;D_1__Actinobacteria;D_2__Thermoleophilia;D_3__Solirubrobacterales;D_4__67-14;D_5__metagenome |
| D_0__Bacteria;D_1__Firmicutes;D_2__Clostridia;D_3__Clostridiales;D_4__Lachnospiraceae;D_5__Acetitomaculum |
| D_0__Bacteria;D_1__Firmicutes;D_2__Bacilli;D_3__Bacillales;D_4__Bacillaceae;D_5__Bacillus |
| D_0__Bacteria;D_1__Firmicutes;D_2__Clostridia;D_3__Clostridiales;D_4__Ruminococcaceae;D_5__uncultured |
| D_0__Bacteria;D_1__Actinobacteria;D_2__Coriobacteriia;D_3__Coriobacteriales;D_4__Atopobiaceae;D_5__Atopobium |

| no diet filtering | | diet filtering | |
|---|---|---|---|
| clr1 | W1 | clr | W |
| **-7.177184652** | **237** | NA | NA |
| **-4.505135111** | **235** | **4.509352998** | **231** |
| **-4.549212311** | **235** | **4.553430198** | **231** |
| **-4.831113346** | **235** | **4.835331232** | **231** |
| **5.417597145** | **234** | NA | NA |
| **4.7560705** | **234** | **-4.751852614** | **231** |
| **3.727956306** | **227** | **-3.72373842** | **225** |
| **-3.805742911** | **227** | **3.809960797** | **223** |
| **-3.485964285** | **217** | **3.490182172** | **213** |
| **-3.570998273** | **216** | **3.575216159** | **212** |
| **-3.171359546** | **208** | **3.175577432** | **204** |
| **-3.065866252** | **204** | **3.070084138** | **200** |
| **-3.10950574** | **201** | **3.113723627** | **198** |
| **-2.926927869** | **200** | **2.931145755** | **198** |
| **-3.275241127** | **200** | **3.279459014** | **196** |
| **-2.877343419** | **197** | **2.881561305** | **194** |
| **-3.039221892** | **196** | **3.043439778** | **193** |
| **-2.641291564** | **190** | **2.64550945** | **188** |
| **-2.713238632** | **190** | **2.717456518** | **188** |
| **-2.635427393** | **189** | **2.639645279** | **186** |
| -2.317852973 | 174 | 2.322070859 | 171 |
| -2.001562971 | 164 | 2.005780857 | 161 |
| -2.112338079 | 162 | 2.116555965 | 159 |
| -1.612544827 | 154 | 1.616762713 | 151 |
| -2.304256762 | 152 | 2.308474649 | 150 |
| -2.089944586 | 151 | 2.094162472 | 148 |
| -1.32901941 | 138 | 1.333237297 | 135 |

D_0__Bacteria;D_1__Proteobacteria;D_2__Gammaproteobacteria;D_3__Aeromonadales;D_4__Aeromonadaceae;D_5__Aeromonas
D_0__Bacteria;D_1__Firmicutes;D_2__Bacilli;D_3__Lactobacillales;D_4__Lactobacillaceae;D_5__Lactobacillus
D_0__Bacteria;D_1__Firmicutes;D_2__Clostridia;D_3__Clostridiales;D_4__TC1;D_5__uncultured Firmicutes bacterium
D_0__Bacteria;D_1__Actinobacteria;D_2__Actinobacteria;D_3__Corynebacteriales;D_4__Nocardiaceae;D_5__Millisia
D_0__Bacteria;D_1__Proteobacteria;D_2__Gammaproteobacteria;D_3__Betaproteobacteriales;D_4__Burkholderiaceae;D_5__Hydrogenophaga
D_0__Bacteria;D_1__Actinobacteria;D_2__Acidimicrobiia;D_3__Microtrichales;D_4__Microtrichaceae;D_5__IMCC26207
D_0__Bacteria;D_1__Actinobacteria;D_2__Actinobacteria;D_3__Corynebacteriales;D_4__Nocardiaceae;D_5__Rhodococcus
D_0__Bacteria;D_1__Proteobacteria;D_2__Gammaproteobacteria;D_3__Legionellales;D_4__Legionellaceae;D_5__Legionella
D_0__Bacteria;D_1__Proteobacteria;D_2__Gammaproteobacteria;D_3__Gammaproteobacteria Incertae Sedis;D_4__Unknown Family;Ambiguous_taxa
D_0__Bacteria;D_1__Actinobacteria;D_2__Actinobacteria;D_3__Micrococcales;D_4__Micrococcaceae;D_5__Kocuria
D_0__Bacteria;D_1__Chloroflexi;D_2__Anaerolineae;D_3__Caldilineales;D_4__Caldilineaceae;D_5__uncultured
D_0__Bacteria;D_1__Actinobacteria;D_2__Actinobacteria;D_3__Micrococcales;D_4__Brevibacteriaceae;D_5__Brevibacterium
D_0__Bacteria;D_1__Proteobacteria;D_2__Gammaproteobacteria;__;__;__
D_0__Bacteria;D_1__Actinobacteria;D_2__Actinobacteria;D_3__Micrococcales;D_4__Microbacteriaceae;D_5__Microbacterium
D_0__Bacteria;D_1__Proteobacteria;D_2__Alphaproteobacteria;D_3__Rhizobiales;D_4__Rhizobiaceae;D_5__Hoeflea
D_0__Bacteria;D_1__Proteobacteria;D_2__Gammaproteobacteria;D_3__Betaproteobacteriales;D_4__Burkholderiaceae;D_5__Undibacterium
D_0__Bacteria;D_1__Proteobacteria;D_2__Alphaproteobacteria;D_3__Rhodobacterales;D_4__Rhodobacteraceae;D_5__Paracoccus
D_0__Bacteria;D_1__Chlamydiae;D_2__Chlamydiae;D_3__Chlamydiales;D_4__Parachlamydiaceae;D_5__uncultured
D_0__Bacteria;D_1__Proteobacteria;D_2__Alphaproteobacteria;D_3__Rhodobacterales;D_4__Rhodobacteraceae;D_5__Amaricoccus
D_0__Bacteria;D_1__Bacteroidetes;D_2__Bacteroidia;D_3__Cytophagales;D_4__Cyclobacteriaceae;D_5__Roseivirga
D_0__Bacteria;D_1__Proteobacteria;D_2__Alphaproteobacteria;D_3__Paracaedibacterales;D_4__Paracaedibacteraceae;D_5__uncultured
D_0__Bacteria;D_1__Actinobacteria;D_2__Actinobacteria;D_3__Corynebacteriales;D_4__Nocardiaceae;D_5__Nocardia
D_0__Bacteria;D_1__Proteobacteria;D_2__Alphaproteobacteria;D_3__Rhodobacterales;D_4__Rhodobacteraceae;__
D_0__Bacteria;D_1__Proteobacteria;D_2__Gammaproteobacteria;D_3__Pseudomonadales;D_4__Moraxellaceae;D_5__Acinetobacter
D_0__Bacteria;D_1__Actinobacteria;D_2__Actinobacteria;D_3__Micrococcales;D_4__Dermacoccaceae;D_5__Dermacoccus
D_0__Bacteria;D_1__Proteobacteria;D_2__Gammaproteobacteria;D_3__Oceanospirillales;D_4__Pseudohongiellaceae;D_5__Pseudohongiella
D_0__Bacteria;D_1__Actinobacteria;D_2__Acidimicrobiia;D_3__Microtrichales;D_4__Microtrichaceae;D_5__Candidatus Microthrix
D_0__Bacteria;D_1__Proteobacteria;D_2__Alphaproteobacteria;D_3__Rhodospirillales;D_4__Terasakiellaceae;D_5__uncultured
D_0__Bacteria;D_1__Proteobacteria;D_2__Gammaproteobacteria;D_3__Ga0077536;Ambiguous_taxa;Ambiguous_taxa
D_0__Bacteria;D_1__Proteobacteria;D_2__Alphaproteobacteria;D_3__Sphingomonadales;D_4__Sphingomonadaceae;D_5__Sphingomonas
D_0__Bacteria;D_1__Bacteroidetes;D_2__Bacteroidia;D_3__Flavobacteriales;D_4__Flavobacteriaceae;D_5__Mesonia
D_0__Bacteria;D_1__Proteobacteria;D_2__Alphaproteobacteria;D_3__Rhizobiales;D_4__Rhizobiaceae;D_5__Allorhizobium-Neorhizobium-Pararhizobium-Rhizobium
D_0__Bacteria;D_1__Proteobacteria;D_2__Alphaproteobacteria;D_3__Rhizobiales;D_4__Stappiaceae;D_5__Pannonibacter

| | | | | |
|---|---|---|---|---|
| 3.563957462 | 127 | | -3.559739575 | 126 |
| -1.778280252 | 124 | | 1.782498138 | 122 |
| -1.859800197 | 118 | | 1.864018083 | 115 |
| 2.309704684 | 76 | | -2.305486798 | 78 |
| -1.851765003 | 73 | | 1.855982889 | 69 |
| -1.289091885 | 56 | | 1.293309772 | 53 |
| 2.302792038 | 45 | | -2.298574152 | 44 |
| 1.670216626 | 44 | | -1.66599874 | 42 |
| 1.54162491 | 44 | | -1.537407024 | 42 |
| 1.537402751 | 44 | | -1.533184865 | 42 |
| -1.519253445 | 44 | | 1.523471332 | 41 |
| 1.225238645 | 41 | | -1.221020758 | 39 |
| 1.164943147 | 41 | | -1.16072526 | 39 |
| 1.109493366 | 36 | | -1.105275479 | 34 |
| 1.068946855 | 36 | | -1.064728969 | 34 |
| 2.120206947 | 35 | | -2.115989061 | 34 |
| 1.196787395 | 35 | | -1.192569508 | 33 |
| 1.037101482 | 35 | | -1.032883595 | 33 |
| 0.999632137 | 35 | | -0.995414251 | 33 |
| 0.967786764 | 35 | NA | | NA |
| 1.314628909 | 34 | | -1.310411022 | 32 |
| 1.023748342 | 34 | | -1.019530456 | 33 |
| 0.920786401 | 34 | | -0.916568514 | 32 |
| 1.327610003 | 33 | | -1.323392117 | 31 |
| 0.939018556 | 33 | | -0.93480067 | 31 |
| 0.939018556 | 33 | | -0.93480067 | 31 |
| -0.799599382 | 33 | | 0.803817268 | 30 |
| 0.898472046 | 32 | | -0.894254159 | 30 |
| 0.869703838 | 32 | | -0.865485952 | 30 |
| 0.796463019 | 32 | | -0.792245133 | 30 |
| 0.713632797 | 32 | | -0.709414911 | 30 |
| 1.042650667 | 31 | | -1.03843278 | 29 |
| 0.908475532 | 31 | | -0.904257646 | 29 |

D_0__Bacteria;D_1__Planctomycetes;D_2__Planctomycetacia;D_3__Pirellulales;D_4__Pirellulaceae;D_5__Pirellula
D_0__Bacteria;D_1__Proteobacteria;D_2__Alphaproteobacteria;D_3__Caulobacterales;D_4__Caulobacteraceae;D_5__Caulobacter
D_0__Bacteria;D_1__Bacteroidetes;D_2__Bacteroidia;D_3__Flavobacteriales;D_4__Flavobacteriaceae;D_5__Maribacter
D_0__Bacteria;D_1__Bacteroidetes;D_2__Bacteroidia;D_3__Flavobacteriales;D_4__Cryomorphaceae;D_5__Owenweeksia
D_0__Bacteria;D_1__Proteobacteria;D_2__Alphaproteobacteria;D_3__Sphingomonadales;D_4__Sphingomonadaceae;D_5__Novosphingobium
D_0__Bacteria;D_1__Proteobacteria;D_2__Gammaproteobacteria;D_3__Pasteurellales;D_4__Pasteurellaceae;D_5__Haemophilus
D_0__Bacteria;D_1__Firmicutes;D_2__Clostridia;D_3__Clostridiales;D_4__Lachnospiraceae;D_5__Vallitalea
D_0__Bacteria;D_1__Actinobacteria;D_2__Acidimicrobiia;D_3__Microtrichales;D_4__Iamiaceae;D_5__Iamia
D_0__Bacteria;D_1__Proteobacteria;D_2__Gammaproteobacteria;D_3__Oceanospirillales;D_4__Halomonadaceae;D_5__Halomonas
D_0__Bacteria;D_1__Bacteroidetes;D_2__Bacteroidia;D_3__Cytophagales;D_4__Microscillaceae;D_5__uncultured
D_0__Bacteria;D_1__Planctomycetes;D_2__Planctomycetacia;D_3__Gemmatales;D_4__Gemmataceae;D_5__Gemmata
D_0__Bacteria;D_1__Verrucomicrobia;D_2__Verrucomicrobiae;D_3__Verrucomicrobiales;D_4__Verrucomicrobiaceae;D_5__Prosthecobacter
D_0__Bacteria;D_1__Bacteroidetes;D_2__Bacteroidia;D_3__Flavobacteriales;D_4__Flavobacteriaceae;D_5__Cellulophaga
D_0__Bacteria;D_1__Bacteroidetes;D_2__Bacteroidia;D_3__Sphingobacteriales;D_4__NS11-12 marine group;D_5__uncultured bacterium
D_0__Bacteria;D_1__Bacteroidetes;D_2__Bacteroidia;D_3__Flavobacteriales;D_4__Flavobacteriaceae;D_5__Arenibacter
D_0__Bacteria;D_1__Proteobacteria;D_2__Alphaproteobacteria;D_3__Holosporales;D_4__Holosporaceae;D_5__uncultured
D_0__Bacteria;D_1__Proteobacteria;D_2__Alphaproteobacteria;D_3__Rhodospirillales;D_4__Rhodospirillaceae;D_5__uncultured
D_0__Bacteria;D_1__Deinococcus-Thermus;D_2__Deinococci;D_3__Thermales;D_4__Thermaceae;D_5__Thermus
D_0__Bacteria;D_1__Actinobacteria;D_2__Actinobacteria;D_3__Frankiales;D_4__Nakamurellaceae;D_5__Nakamurella
D_0__Bacteria;D_1__Firmicutes;D_2__Bacilli;D_3__Lactobacillales;D_4__Leuconostocaceae;D_5__Weissella
D_0__Bacteria;D_1__Proteobacteria;D_2__Alphaproteobacteria;D_3__Rhodobacterales;D_4__Rhodobacteraceae;D_5__Gemmobacter
D_0__Bacteria;D_1__Bacteroidetes;D_2__Bacteroidia;D_3__Flavobacteriales;D_4__Crocinitomicaceae;D_5__Fluviicola
D_0__Bacteria;D_1__Actinobacteria;D_2__Actinobacteria;D_3__Pseudonocardiales;D_4__Pseudonocardiaceae;D_5__Pseudonocardia
D_0__Bacteria;D_1__Bacteroidetes;D_2__Bacteroidia;D_3__Cytophagales;D_4__Spirosomaceae;__
D_0__Bacteria;D_1__Proteobacteria;D_2__Gammaproteobacteria;D_3__Pseudomonadales;D_4__Moraxellaceae;D_5__Enhydrobacter
D_0__Bacteria;D_1__Actinobacteria;D_2__Thermoleophilia;D_3__Solirubrobacterales;D_4__67-14;__
D_0__Bacteria;D_1__Bacteroidetes;D_2__Bacteroidia;D_3__Cytophagales;D_4__Spirosomaceae;D_5__Taeseokella
D_0__Bacteria;D_1__Proteobacteria;D_2__Alphaproteobacteria;D_3__Rhizobiales;D_4__Beijerinckiaceae;D_5__Bosea
D_0__Bacteria;D_1__Planctomycetes;D_2__Planctomycetacia;D_3__Isosphaerales;D_4__Isosphaeraceae;D_5__uncultured
D_0__Bacteria;D_1__Proteobacteria;D_2__Gammaproteobacteria;D_3__Salinisphaerales;D_4__Solimonadaceae;D_5__Nevskia
D_0__Bacteria;D_1__Planctomycetes;D_2__Planctomycetacia;D_3__Planctomycetales;D_4__Gimesiaceae;D_5__uncultured
D_0__Bacteria;D_1__Planctomycetes;D_2__OM190;__;__;__
D_0__Bacteria;D_1__Actinobacteria;D_2__KIST-JJY010;D_3__uncultured bacterium;D_4__uncultured bacterium;D_5__uncultured bacterium

| | | | |
|---|---|---|---|
| 0.713632797 | 31 | -0.709414911 | 29 |
| 0.706194679 | 31 | -0.701976793 | 29 |
| 0.700117294 | 31 | -0.695899408 | 29 |
| 0.694978938 | 31 | -0.690761052 | 29 |
| 0.690527891 | 31 | -0.686310005 | 29 |
| 0.658358528 | 31 | -0.654140642 | 29 |
| 0.646469363 | 31 | -0.642251477 | 29 |
| 0.586011418 | 31 | -0.581793531 | 29 |
| 0.547972354 | 31 | -0.543754468 | 29 |
| 0.713632797 | 30 | -0.709414911 | 28 |
| 0.713632797 | 30 | -0.709414911 | 28 |
| 0.713632797 | 30 | -0.709414911 | 28 |
| 0.700117294 | 30 | -0.695899408 | 28 |
| 0.700117294 | 30 | -0.695899408 | 28 |
| 0.671874032 | 30 | -0.667656146 | 28 |
| 0.653907482 | 30 | -0.649689596 | 28 |
| 0.646469363 | 30 | -0.642251477 | 28 |
| 0.644318079 | 30 | -0.640100193 | 28 |
| 0.568504272 | 30 | -0.564286386 | 29 |
| 0.501994831 | 30 | -0.497776945 | 29 |
| 1.212916501 | 29 | -1.208698615 | 28 |
| 0.88573217 | 29 | -0.881514284 | 27 |
| 0.864040537 | 29 | -0.859822651 | 27 |
| 0.7232222 | 29 | -0.719004314 | 27 |
| 0.713632797 | 29 | -0.709414911 | 27 |
| 0.669574269 | 29 | -0.665356383 | 28 |
| 0.6613456 | 29 | -0.657127714 | 27 |
| 0.653907482 | 29 | -0.649689596 | 27 |
| 0.648769126 | 29 | -0.64455124 | 27 |
| 0.641331007 | 29 | -0.637113121 | 27 |
| 0.63295386 | 29 | -0.628735974 | 27 |
| 0.623364457 | 29 | -0.619146571 | 27 |
| 0.619192353 | 29 | -0.614974466 | 27 |

D_0__Bacteria;D_1__Proteobacteria;D_2__Alphaproteobacteria;D_3__Caulobacterales;D_4__Hyphomonadaceae;D_5__Hyphomonas
D_0__Bacteria;D_1__Actinobacteria;D_2__Actinobacteria;D_3__Propionibacteriales;D_4__Nocardioidaceae;D_5__Nocardioides
D_0__Bacteria;D_1__Chloroflexi;D_2__Anaerolineae;D_3__SBR1031;D_4__uncultured bacterium;D_5__uncultured bacterium
D_0__Bacteria;D_1__Proteobacteria;D_2__Alphaproteobacteria;D_3__Sphingomonadales;D_4__Sphingomonadaceae;D_5__Sphingopyxis
D_0__Bacteria;D_1__Chlamydiae;D_2__Chlamydiae;D_3__Chlamydiales;D_4__Criblamydiaceae;__
D_0__Bacteria;D_1__Firmicutes;D_2__Clostridia;D_3__Clostridiales;D_4__Syntrophomonadaceae;D_5__Syntrophomonas
D_0__Bacteria;D_1__Proteobacteria;D_2__Alphaproteobacteria;D_3__Rhizobiales;D_4__Beijerinckiaceae;__
D_0__Bacteria;D_1__Actinobacteria;D_2__Actinobacteria;D_3__Micrococcales;D_4__Dermatophilaceae;D_5__Kineosphaera
D_0__Bacteria;D_1__Patescibacteria;D_2__Saccharimonadia;D_3__Saccharimonadales;D_4__uncultured bacterium;D_5__uncultured bacterium
D_0__Bacteria;D_1__Actinobacteria;D_2__Actinobacteria;D_3__PeM15;D_4__uncultured soil bacterium;D_5__uncultured soil bacterium
D_0__Bacteria;D_1__Firmicutes;D_2__Bacilli;D_3__Lactobacillales;D_4__Aerococcaceae;D_5__Dolosicoccus
D_0__Bacteria;D_1__Firmicutes;D_2__Clostridia;D_3__Clostridiales;D_4__Lachnospiraceae;D_5__Coprococcus 2
D_0__Bacteria;D_1__Bacteroidetes;D_2__Bacteroidia;D_3__Cytophagales;D_4__Cyclobacteriaceae;D_5__uncultured
D_0__Bacteria;D_1__Proteobacteria;D_2__Deltaproteobacteria;D_3__Myxococcales;D_4__Phaselicystidaceae;D_5__Phaselicystis
D_0__Bacteria;D_1__Proteobacteria;D_2__Alphaproteobacteria;D_3__Rhizobiales;D_4__Devosiaceae;D_5__uncultured
D_0__Bacteria;D_1__Proteobacteria;D_2__Alphaproteobacteria;D_3__Micavibrionales;D_4__Micavibrionaceae;D_5__uncultured
D_0__Bacteria;D_1__Firmicutes;D_2__Clostridia;D_3__Clostridiales;D_4__Lachnospiraceae;D_5__[Eubacterium] ruminantium group
D_0__Bacteria;D_1__Proteobacteria;D_2__Gammaproteobacteria;D_3__Betaproteobacteriales;D_4__Nitrosomonadaceae;D_5__oc32
D_0__Bacteria;D_1__Verrucomicrobia;D_2__Verrucomicrobiae;D_3__Pedosphaerales;D_4__Pedosphaeraceae;D_5__uncultured verrucomicrobium DEV059
D_0__Bacteria;D_1__Proteobacteria;D_2__Alphaproteobacteria;D_3__Caulobacterales;D_4__Hyphomonadaceae;D_5__Hirschia
D_0__Bacteria;D_1__Proteobacteria;D_2__Gammaproteobacteria;D_3__Pseudomonadales;D_4__Moraxellaceae;D_5__Psychrobacter
D_0__Bacteria;D_1__Actinobacteria;D_2__Coriobacteriia;D_3__Coriobacteriales;D_4__Eggerthellaceae;D_5__Gordonibacter
D_0__Bacteria;D_1__Proteobacteria;D_2__Gammaproteobacteria;D_3__Alteromonadales;D_4__Marinobacteraceae;D_5__Marinobacter
D_0__Bacteria;D_1__Proteobacteria;D_2__Alphaproteobacteria;D_3__Rhodobacterales;D_4__Rhodobacteraceae;D_5__Defluviimonas
D_0__Bacteria;D_1__Planctomycetes;D_2__Planctomycetacia;D_3__Planctomycetales;D_4__Rubinisphaeraceae;D_5__SH-PL14
D_0__Bacteria;D_1__Proteobacteria;D_2__Alphaproteobacteria;D_3__Caulobacterales;D_4__Caulobacteraceae;D_5__Brevundimonas
D_0__Bacteria;D_1__Bacteroidetes;D_2__Bacteroidia;D_3__Cytophagales;D_4__Microscillaceae;D_5__OLB12
D_0__Bacteria;D_1__Firmicutes;D_2__Clostridia;D_3__Clostridiales;D_4__Lachnospiraceae;D_5__Epulopiscium
D_0__Bacteria;D_1__Firmicutes;D_2__Bacilli;D_3__Bacillales;D_4__Alicyclobacillaceae;D_5__Tumebacillus
D_0__Bacteria;D_1__Firmicutes;D_2__Clostridia;D_3__Clostridiales;D_4__Eubacteriaceae;D_5__Acetobacterium
D_0__Bacteria;D_1__Firmicutes;D_2__Bacilli;D_3__Lactobacillales;D_4__Leuconostocaceae;D_5__Leuconostoc
D_0__Bacteria;D_1__Firmicutes;D_2__Clostridia;D_3__Clostridiales;D_4__Lachnospiraceae;D_5__Lachnospiraceae UCG-010
D_0__Bacteria;D_1__Firmicutes;D_2__Clostridia;D_3__Clostridiales;D_4__Peptostreptococcaceae;D_5__Paeniclostridium

| | | | |
|---|---|---|---|
| 0.617287072 | 29 | -0.613069186 | 27 |
| 0.615926339 | 29 | -0.611708453 | 27 |
| 0.609848954 | 29 | -0.605631068 | 28 |
| 0.579305929 | 29 | -0.575088043 | 27 |
| 0.565938904 | 29 | -0.561721018 | 27 |
| 0.554893666 | 29 | -0.55067578 | 27 |
| 0.554328681 | 29 | -0.550110795 | 27 |
| 0.545951533 | 29 | -0.541733647 | 27 |
| 0.543434615 | 29 | -0.539216729 | 27 |
| 0.503667823 | 29 | -0.499449936 | 28 |
| 0.479783198 | 29 | -0.475565312 | 27 |
| 0.399883979 | 29 | -0.395666093 | 27 |
| 0.659984867 | 28 | -0.655766981 | 26 |
| 0.658358528 | 28 | -0.654140642 | 26 |
| 0.652546749 | 28 | -0.648328862 | 26 |
| 0.646469363 | 28 | -0.642251477 | 27 |
| 0.629441843 | 28 | -0.625223956 | 26 |
| 0.598633213 | 28 | -0.594415326 | 26 |
| 0.595121195 | 28 | -0.590903309 | 26 |
| 0.592821433 | 28 | -0.588603547 | 26 |
| 0.59192816 | 28 | -0.587710274 | 26 |
| 0.591514076 | 28 | -0.58729619 | 26 |
| 0.590462464 | 28 | -0.586244578 | 26 |
| 0.576196332 | 28 | -0.571978445 | 26 |
| 0.564388237 | 28 | -0.560170351 | 26 |
| 0.560127125 | 28 | -0.555909238 | 26 |
| 0.558500786 | 28 | -0.5542829 | 26 |
| 0.538907897 | 28 | -0.534690011 | 26 |
| 0.530944833 | 28 | -0.526726947 | 26 |
| 0.488160346 | 28 | -0.48394246 | 26 |
| 0.479182581 | 28 | -0.474964695 | 26 |
| 0.457016705 | 28 | -0.452798818 | 26 |
| 0.433286558 | 28 | -0.429068672 | 26 |

D_0__Bacteria;D_1__Proteobacteria;D_2__Deltaproteobacteria;D_3__Syntrophobacterales;D_4__Syntrophaceae;D_5__Desulfomonile

D_0__Bacteria;D_1__Chlamydiae;D_2__Chlamydiae;D_3__Chlamydiales;D_4__Parachlamydiaceae;__

D_0__Bacteria;D_1__Chloroflexi;D_2__Chloroflexia;D_3__Thermomicrobiales;D_4__JG30-KF-CM45;D_5__uncultured bacterium

D_0__Bacteria;D_1__Firmicutes;D_2__Clostridia;D_3__Clostridiales;D_4__Lachnospiraceae;D_5__Anaerostipes

D_0__Bacteria;D_1__Actinobacteria;D_2__Coriobacteriia;D_3__Coriobacteriales;D_4__Eggerthellaceae;D_5__Eggerthella

D_0__Bacteria;D_1__Planctomycetes;D_2__Planctomycetacia;D_3__Gemmatales;D_4__Gemmataceae;D_5__Fimbriiglobus

D_0__Bacteria;D_1__Actinobacteria;D_2__Coriobacteriia;D_3__Coriobacteriales;D_4__Eggerthellaceae;D_5__Senegalimassilia

D_0__Bacteria;D_1__Firmicutes;D_2__Clostridia;D_3__Clostridiales;D_4__Lachnospiraceae;D_5__Cellulosilyticum

D_0__Bacteria;D_1__Actinobacteria;D_2__Coriobacteriia;D_3__Coriobacteriales;D_4__uncultured;D_5__uncultured Coriobacteriaceae bacterium

D_0__Bacteria;D_1__Patescibacteria;D_2__Saccharimonadia;D_3__Saccharimonadales;D_4__wastewater metagenome;D_5__wastewater metagenome

D_0__Bacteria;D_1__Firmicutes;D_2__Clostridia;D_3__Clostridiales;D_4__Family XIII;D_5__Family XIII AD3011 group

D_0__Bacteria;D_1__Proteobacteria;D_2__Alphaproteobacteria;D_3__Rickettsiales;D_4__Rickettsiaceae;D_5__uncultured

D_0__Bacteria;D_1__Actinobacteria;D_2__Acidimicrobiia;D_3__Microtrichales;D_4__uncultured;D_5__uncultured bacterium

D_0__Bacteria;D_1__Proteobacteria;D_2__Alphaproteobacteria;D_3__Rhodospirillales;D_4__Rhodospirillaceae;D_5__Haematospirillum

D_0__Bacteria;D_1__Proteobacteria;D_2__Gammaproteobacteria;D_3__Betaproteobacteriales;D_4__Methylophilaceae;D_5__Methylotenera

D_0__Bacteria;D_1__Proteobacteria;D_2__Gammaproteobacteria;D_3__Oceanospirillales;D_4__Alcanivoracaceae;D_5__Alcanivorax

D_0__Bacteria;D_1__Firmicutes;D_2__Clostridia;D_3__Clostridiales;D_4__Lachnospiraceae;D_5__Roseburia

D_0__Bacteria;D_1__Proteobacteria;D_2__Alphaproteobacteria;D_3__Rhizobiales;D_4__Rhizobiales Incertae Sedis;D_5__Bauldia

D_0__Bacteria;D_1__Proteobacteria;D_2__Gammaproteobacteria;D_3__Pseudomonadales;D_4__Moraxellaceae;D_5__Alkanindiges

D_0__Bacteria;D_1__Proteobacteria;D_2__Alphaproteobacteria;D_3__Reyranellales;D_4__Reyranellaceae;D_5__Reyranella

D_0__Bacteria;D_1__Chlamydiae;D_2__Chlamydiae;D_3__Chlamydiales;D_4__Parachlamydiaceae;D_5__Candidatus Protochlamydia

D_0__Bacteria;D_1__Planctomycetes;D_2__Planctomycetacia;D_3__Pirellulales;D_4__Pirellulaceae;D_5__Pir4 lineage

D_0__Bacteria;D_1__Actinobacteria;D_2__Acidimicrobiia;D_3__IMCC26256;__;__

D_0__Bacteria;D_1__Proteobacteria;D_2__Alphaproteobacteria;D_3__Rhizobiales;__;__

D_0__Bacteria;D_1__Actinobacteria;D_2__Coriobacteriia;D_3__Coriobacteriales;D_4__Coriobacteriaceae;D_5__Collinsella

D_0__Bacteria;D_1__Firmicutes;D_2__Erysipelotrichia;D_3__Erysipelotrichales;D_4__Erysipelotrichaceae;D_5__Breznakia

D_0__Bacteria;D_1__Firmicutes;D_2__Clostridia;D_3__Clostridiales;D_4__Family XI;D_5__Gottschalkia

D_0__Bacteria;D_1__Bacteroidetes;D_2__Bacteroidia;D_3__Flavobacteriales;D_4__Flavobacteriaceae;D_5__Flavobacterium

D_0__Bacteria;D_1__Proteobacteria;D_2__Deltaproteobacteria;D_3__Myxococcales;D_4__Nannocystaceae;D_5__Nannocystis

D_0__Bacteria;D_1__Proteobacteria;D_2__Gammaproteobacteria;D_3__Pseudomonadales;D_4__Moraxellaceae;D_5__Perlucidibaca

D_0__Bacteria;D_1__Proteobacteria;D_2__Alphaproteobacteria;D_3__Rhizobiales;D_4__Hyphomicrobiaceae;__

D_0__Bacteria;D_1__Proteobacteria;D_2__Gammaproteobacteria;D_3__Salinisphaerales;D_4__Solimonadaceae;D_5__Polycyclovorans

D_0__Bacteria;D_1__Actinobacteria;D_2__Actinobacteria;D_3__PeM15;__;__

| | | | |
|---|---|---|---|
| 0.433205058 | 28 | -0.428987172 | 27 |
| 0.422690659 | 28 | -0.418472772 | 26 |
| 0.4225627 | 28 | -0.418344813 | 26 |
| 0.405416817 | 28 | -0.401198931 | 26 |
| 0.329470996 | 28 | -0.32525311 | 26 |
| 0.328147174 | 28 | -0.323929288 | 26 |
| 0.280216192 | 28 | -0.275998305 | 26 |
| 0.276430678 | 28 | -0.272212792 | 26 |
| 0.201012421 | 28 | -0.196794535 | 26 |
| 0.16750034 | 28 | -0.163282453 | 26 |
| 0.113329968 | 28 | -0.109112082 | 26 |
| 0.952925095 | 27 | -0.948707209 | 25 |
| 0.609583348 | 27 | -0.605365462 | 25 |
| 0.599436464 | 27 | -0.595218578 | 25 |
| 0.588356501 | 27 | -0.584138614 | 25 |
| 0.565938904 | 27 | -0.561721018 | 25 |
| 0.560127125 | 27 | -0.555909238 | 25 |
| 0.540534236 | 27 | -0.53631635 | 25 |
| 0.533096117 | 27 | -0.528878231 | 25 |
| 0.478889925 | 27 | -0.474672039 | 25 |
| 0.430196322 | 27 | -0.425978436 | 25 |
| 0.285213911 | 27 | -0.280996024 | 25 |
| 0.186435834 | 27 | -0.182217948 | 25 |
| 0.17606402 | 27 | -0.171846134 | 25 |
| -0.941905659 | 27 | 0.946123545 | 25 |
| -1.616728841 | 27 | 1.620946727 | 25 |
| -1.867231661 | 27 | 1.871449547 | 25 |
| 1.283012594 | 26 | -1.278794708 | 24 |
| 0.602559314 | 26 | -0.598341428 | 24 |
| 0.495168351 | 26 | -0.490950464 | 24 |
| 0.475630905 | 26 | -0.471413019 | 24 |
| 0.399029024 | 26 | -0.394811138 | 24 |
| 0.290613465 | 26 | -0.286395579 | 24 |

D_0__Bacteria;D_1__Firmicutes;D_2__Clostridia;D_3__Clostridiales;D_4__Lachnospiraceae;D_5__Lachnospiraceae ND3007 group
D_0__Bacteria;D_1__Patescibacteria;D_2__Saccharimonadia;D_3__Saccharimonadales;D_4__uncultured candidate division WS5 bacterium;D_5__uncultured candidate division WS5 bacterium
D_0__Bacteria;D_1__Proteobacteria;D_2__Alphaproteobacteria;D_3__Rhizobiales;D_4__Devosiaceae;D_5__Devosia
D_0__Bacteria;D_1__Actinobacteria;D_2__Actinobacteria;D_3__Propionibacteriales;D_4__Propionibacteriaceae;D_5__uncultured
D_0__Bacteria;D_1__Planctomycetes;D_2__Planctomycetacia;D_3__Pirellulales;D_4__Pirellulaceae;D_5__Blastopirellula
D_0__Bacteria;D_1__Actinobacteria;D_2__Acidimicrobiia;D_3__Microtrichales;D_4__Ilumatobacteraceae;D_5__CL500-29 marine group
D_0__Bacteria;D_1__Firmicutes;D_2__Bacilli;D_3__Lactobacillales;D_4__Carnobacteriaceae;D_5__Trichococcus
D_0__Bacteria;D_1__Proteobacteria;D_2__Alphaproteobacteria;D_3__Rhizobiales;D_4__Beijerinckiaceae;D_5__uncultured
D_0__Bacteria;D_1__Firmicutes;D_2__Clostridia;D_3__Clostridiales;D_4__Family XI;D_5__Gallicola
D_0__Bacteria;D_1__Firmicutes;D_2__Clostridia;D_3__Clostridiales;D_4__Ruminococcaceae;D_5__Ruminococcus 2
D_0__Bacteria;D_1__Planctomycetes;D_2__Planctomycetacia;D_3__Pirellulales;D_4__Pirellulaceae;D_5__uncultured
D_0__Bacteria;D_1__Fusobacteria;D_2__Fusobacteriia;D_3__Fusobacteriales;D_4__Fusobacteriaceae;D_5__Cetobacterium
D_0__Bacteria;D_1__Firmicutes;D_2__Clostridia;D_3__Clostridiales;D_4__Clostridiaceae 1;D_5__Clostridium sensu stricto 13
D_0__Bacteria;D_1__Chloroflexi;D_2__Chloroflexia;D_3__Thermomicrobiales;D_4__JG30-KF-CM45;D_5__Paraburkholderia tropica
D_0__Bacteria;D_1__Actinobacteria;D_2__Actinobacteria;D_3__PeM15;D_4__metagenome;D_5__metagenome
D_0__Bacteria;D_1__Firmicutes;D_2__Clostridia;D_3__Clostridiales;D_4__Lachnospiraceae;D_5__[Eubacterium] hallii group
D_0__Bacteria;D_1__Firmicutes;D_2__Clostridia;D_3__Clostridiales;D_4__Ruminococcaceae;D_5__Ruminococcus 1
D_0__Bacteria;D_1__Proteobacteria;D_2__Gammaproteobacteria;D_3__Oceanospirillales;D_4__Saccharospirillaceae;D_5__Oceanobacter
D_0__Bacteria;D_1__Firmicutes;D_2__Bacilli;D_3__Bacillales;D_4__Staphylococcaceae;D_5__Staphylococcus
D_0__Bacteria;D_1__Bacteroidetes;D_2__Bacteroidia;D_3__Flavobacteriales;D_4__Flavobacteriaceae;__
D_0__Bacteria;D_1__Firmicutes;D_2__Clostridia;D_3__Clostridiales;D_4__Lachnospiraceae;D_5__Blautia
D_0__Bacteria;D_1__Chlamydiae;D_2__Chlamydiae;D_3__Chlamydiales;D_4__Parachlamydiaceae;D_5__Neochlamydia
D_0__Bacteria;D_1__Firmicutes;D_2__Clostridia;D_3__Clostridiales;D_4__Eubacteriaceae;__
D_0__Bacteria;D_1__Actinobacteria;D_2__Actinobacteria;D_3__Micrococcales;D_4__Micrococcaceae;D_5__Rothia
D_0__Bacteria;D_1__Proteobacteria;D_2__Gammaproteobacteria;D_3__Betaproteobacteriales;D_4__Rhodocyclaceae;D_5__Zoogloea
D_0__Bacteria;D_1__Actinobacteria;D_2__Actinobacteria;D_3__Micrococcales;D_4__Micrococcaceae;__
D_0__Bacteria;D_1__Firmicutes;D_2__Clostridia;D_3__Clostridiales;D_4__Ruminococcaceae;D_5__Saccharofermentans
D_0__Bacteria;D_1__Firmicutes;D_2__Clostridia;D_3__Clostridiales;D_4__Lachnospiraceae;D_5__[Ruminococcus] torques group
D_0__Bacteria;D_1__Planctomycetes;D_2__Planctomycetacia;D_3__Gemmatales;D_4__Gemmataceae;D_5__uncultured
D_0__Bacteria;D_1__Proteobacteria;D_2__Alphaproteobacteria;D_3__Rhodobacterales;D_4__Rhodobacteraceae;D_5__Pseudorhodobacter
D_0__Bacteria;D_1__Proteobacteria;D_2__Alphaproteobacteria;D_3__Sphingomonadales;D_4__Sphingomonadaceae;D_5__Sphingobium
D_0__Bacteria;D_1__Planctomycetes;D_2__Planctomycetacia;D_3__Pirellulales;D_4__Pirellulaceae;D_5__Rhodopirellula
D_0__Bacteria;D_1__Proteobacteria;D_2__Gammaproteobacteria;D_3__Betaproteobacteriales;D_4__Rhodocyclaceae;D_5__Methyloversatilis

| | | | |
|---|---|---|---|
| 0.238376356 | 26 | -0.23415847 | 24 |
| -0.059922831 | 26 | 0.064140717 | 24 |
| 0.81713403 | 25 | -0.812916144 | 23 |
| 0.455901773 | 25 | -0.451683887 | 23 |
| 0.425853633 | 25 | -0.421635746 | 23 |
| 0.284183748 | 25 | -0.279965862 | 23 |
| 0.15821698 | 25 | -0.153999094 | 23 |
| 0.07551387 | 25 | -0.071295983 | 23 |
| 0.050495964 | 25 | -0.046278077 | 23 |
| -0.252651067 | 25 | 0.256868953 | 23 |
| -0.780672592 | 25 | 0.784890478 | 23 |
| 0.507425843 | 24 | -0.503207957 | 22 |
| 0.383918965 | 24 | -0.379701079 | 22 |
| 0.34175778 | 24 | -0.337539894 | 22 |
| 0.119908739 | 24 | -0.115690853 | 22 |
| 0.057506662 | 24 | -0.053288776 | 22 |
| -0.576976691 | 24 | 0.581194577 | 22 |
| 1.618514277 | 23 | -1.614296391 | 22 |
| 1.064871087 | 23 | -1.0606532 | 21 |
| 0.395257232 | 23 | -0.391039346 | 21 |
| -0.044163781 | 23 | 0.048381667 | 21 |
| -0.193351058 | 23 | 0.197568944 | 21 |
| -0.362424704 | 23 | 0.36664259 | 21 |
| 0.74055768 | 22 | -0.736339794 | 20 |
| 0.597082545 | 22 | -0.592864659 | 20 |
| 0.507770555 | 22 | -0.503552669 | 20 |
| 0.149008355 | 22 | -0.144790469 | 20 |
| -0.026303055 | 22 | 0.030520941 | 20 |
| -0.602681975 | 22 | 0.606899861 | 20 |
| 0.841857229 | 21 | -0.837639342 | 20 |
| 0.461622057 | 21 | -0.457404171 | 19 |
| 0.290663553 | 21 | -0.286445667 | 19 |
| 0.135593181 | 21 | -0.131375295 | 19 |

D_0__Bacteria;D_1__Proteobacteria;D_2__Alphaproteobacteria;D_3__Rhizobiales;D_4__Xanthobacteraceae;__

D_0__Bacteria;D_1__Proteobacteria;D_2__Alphaproteobacteria;D_3__Rhizobiales;D_4__Hyphomicrobiaceae;D_5__Hyphomicrobium

D_0__Bacteria;D_1__Chloroflexi;D_2__KD4-96;D_3__uncultured bacterium;D_4__uncultured bacterium;D_5__uncultured bacterium

D_0__Bacteria;D_1__Planctomycetes;D_2__Planctomycetacia;D_3__Planctomycetales;D_4__Rubinisphaeraceae;D_5__uncultured

D_0__Bacteria;D_1__Chloroflexi;D_2__Chloroflexia;D_3__Thermomicrobiales;D_4__JG30-KF-CM45;D_5__metagenome

D_0__Bacteria;D_1__Actinobacteria;D_2__Actinobacteria;D_3__Corynebacteriales;D_4__Dietziaceae;D_5__Dietzia

D_0__Bacteria;D_1__Actinobacteria;D_2__Actinobacteria;D_3__Corynebacteriales;D_4__Mycobacteriaceae;D_5__Mycobacterium

D_0__Bacteria;D_1__Firmicutes;D_2__Clostridia;D_3__Clostridiales;D_4__Clostridiaceae 1;__

D_0__Bacteria;D_1__Actinobacteria;D_2__Actinobacteria;D_3__Micrococcales;D_4__Promicromonosporaceae;__

D_0__Bacteria;D_1__Actinobacteria;D_2__Acidimicrobiia;D_3__Microtrichales;D_4__uncultured;__

D_0__Bacteria;D_1__Actinobacteria;D_2__Thermoleophilia;D_3__Solirubrobacterales;D_4__67-14;Ambiguous_taxa

D_0__Bacteria;D_1__Actinobacteria;D_2__Actinobacteria;D_3__PeM15;D_4__uncultured bacterium;D_5__uncultured bacterium

D_0__Bacteria;D_1__Proteobacteria;D_2__Gammaproteobacteria;D_3__Betaproteobacteriales;D_4__Burkholderiaceae;D_5__Simplicispira

D_0__Bacteria;D_1__Proteobacteria;D_2__Alphaproteobacteria;__;__;__

D_0__Bacteria;D_1__Firmicutes;D_2__Clostridia;D_3__Clostridiales;D_4__TC1;D_5__endosymbiont 'TC1' of Trimyema compressum

D_0__Bacteria;D_1__Firmicutes;D_2__Bacilli;D_3__Lactobacillales;D_4__Streptococcaceae;D_5__Streptococcus

D_0__Bacteria;D_1__Actinobacteria;D_2__Actinobacteria;D_3__Corynebacteriales;D_4__Nocardiaceae;D_5__Gordonia

D_0__Bacteria;D_1__Firmicutes;D_2__Clostridia;D_3__Clostridiales;D_4__Clostridiaceae 1;D_5__Sarcina

D_0__Bacteria;D_1__Actinobacteria;D_2__Actinobacteria;D_3__Actinomycetales;D_4__Actinomycetaceae;D_5__Actinomyces

D_0__Bacteria;D_1__Planctomycetes;D_2__OM190;Ambiguous_taxa;Ambiguous_taxa;Ambiguous_taxa

D_0__Bacteria;D_1__Proteobacteria;D_2__Deltaproteobacteria;D_3__Bdellovibrionales;D_4__Bdellovibrionaceae;D_5__OM27 clade

D_0__Bacteria;D_1__Actinobacteria;D_2__Actinobacteria;D_3__Micrococcales;D_4__Sanguibacteraceae;D_5__Sanguibacter

D_0__Bacteria;D_1__Actinobacteria;D_2__Actinobacteria;D_3__Frankiales;D_4__Cryptosporangiaceae;D_5__Fodinicola

D_0__Bacteria;D_1__Proteobacteria;D_2__Alphaproteobacteria;D_3__Rhizobiales;D_4__Xanthobacteraceae;D_5__uncultured

D_0__Bacteria;D_1__Proteobacteria;D_2__Alphaproteobacteria;D_3__Rhizobiales;D_4__Rhizobiales Incertae Sedis;D_5__uncultured

D_0__Bacteria;D_1__Chloroflexi;D_2__Chloroflexia;D_3__Thermomicrobiales;D_4__JG30-KF-CM45;Ambiguous_taxa

D_0__Bacteria;D_1__Firmicutes;D_2__Bacilli;D_3__Lactobacillales;D_4__Streptococcaceae;D_5__Lactococcus

D_0__Bacteria;D_1__Actinobacteria;D_2__Actinobacteria;D_3__Corynebacteriales;D_4__Corynebacteriaceae;D_5__Corynebacterium 1

D_0__Bacteria;D_1__Proteobacteria;D_2__Gammaproteobacteria;D_3__Betaproteobacteriales;D_4__Burkholderiaceae;D_5__Limnobacter

D_0__Bacteria;D_1__Proteobacteria;D_2__Alphaproteobacteria;D_3__Rhizobiales;D_4__Rhizobiaceae;D_5__Shinella

D_0__Bacteria;D_1__Firmicutes;D_2__Bacilli;D_3__Lactobacillales;D_4__Enterococcaceae;D_5__Enterococcus

D_0__Bacteria;D_1__Proteobacteria;D_2__Alphaproteobacteria;D_3__Rhizobiales;D_4__Rhizobiaceae;D_5__Mesorhizobium

D_0__Bacteria;D_1__Firmicutes;D_2__Clostridia;D_3__Clostridiales;D_4__Family XIII;D_5__Anaerovorax

| | | | |
|---|---|---|---|
| -0.009594429 | 21 | 0.013812315 | 19 |
| -0.099447994 | 21 | 0.10366588 | 19 |
| -0.397526982 | 21 | 0.401744868 | 19 |
| -0.422829548 | 21 | 0.427047435 | 19 |
| -0.745991333 | 21 | 0.750209219 | 19 |
| 0.311208661 | 20 | -0.306990775 | 18 |
| 0.197795461 | 20 | -0.193577574 | 18 |
| -0.104272623 | 20 | 0.10849051 | 18 |
| -0.521669478 | 20 | 0.525887364 | 18 |
| -0.611863304 | 20 | 0.61608119 | 18 |
| -0.720586384 | 19 | 0.72480427 | 17 |
| -1.40840509 | 19 | 1.412622976 | 17 |
| 0.696320764 | 18 | -0.692102878 | 16 |
| 0.244985913 | 18 | -0.240768027 | 16 |
| -0.031081963 | 18 | 0.03529985 | 16 |
| -0.087392671 | 18 | 0.091610557 | 16 |
| -0.20318289 | 18 | 0.207400776 | 16 |
| -1.261778716 | 18 | 1.265996602 | 16 |
| -1.331710763 | 18 | 1.33592865 | 16 |
| -0.308471182 | 17 | 0.312689068 | 15 |
| -0.476635154 | 17 | 0.48085304 | 15 |
| -0.638949912 | 17 | 0.643167798 | 15 |
| -0.64021276 | 17 | 0.644430647 | 15 |
| -0.352619983 | 16 | 0.356837869 | 14 |
| -0.470433252 | 16 | 0.474651138 | 14 |
| -0.610188883 | 16 | 0.61440677 | 14 |
| -0.703675163 | 16 | 0.707893049 | 14 |
| -1.05232992 | 15 | 1.056547806 | 13 |
| 0.095206904 | 14 | -0.090989018 | 12 |
| -0.011824086 | 14 | 0.016041972 | 12 |
| -0.34663083 | 14 | 0.350848716 | 12 |
| -0.958653177 | 14 | 0.962871064 | 12 |
| -1.229895116 | 14 | 1.234113002 | 12 |

D_0__Bacteria;D_1__Firmicutes;D_2__Clostridia;D_3__Clostridiales;D_4__Family XIII;__

D_0__Bacteria;D_1__Proteobacteria;D_2__Gammaproteobacteria;D_3__Betaproteobacteriales;D_4__Chitinibacteraceae;D_5__Deefgea

D_0__Bacteria;D_1__Chloroflexi;D_2__Chloroflexia;D_3__Thermomicrobiales;D_4__JG30-KF-CM45;__

D_0__Bacteria;D_1__Proteobacteria;D_2__Gammaproteobacteria;D_3__Betaproteobacteriales;D_4__Burkholderiaceae;__

D_0__Bacteria;D_1__Proteobacteria;D_2__Gammaproteobacteria;D_3__Alteromonadales;D_4__Alteromonadaceae;D_5__Rheinheimera

D_0__Bacteria;D_1__Bacteroidetes;D_2__Bacteroidia;D_3__Cytophagales;D_4__Amoebophilaceae;D_5__Candidatus Amoebophilus

D_0__Bacteria;D_1__Proteobacteria;D_2__Gammaproteobacteria;D_3__Betaproteobacteriales;D_4__Chitinibacteraceae;D_5__Chitinibacter

D_0__Bacteria;D_1__Proteobacteria;D_2__Gammaproteobacteria;D_3__Alteromonadales;D_4__Alteromonadaceae;__

D_0__Bacteria;D_1__Bacteroidetes;D_2__Bacteroidia;D_3__Chitinophagales;D_4__Saprospiraceae;D_5__uncultured

D_0__Bacteria;D_1__Proteobacteria;D_2__Gammaproteobacteria;D_3__Cellvibrionales;D_4__Cellvibrionaceae;D_5__Cellvibrio

D_0__Bacteria;D_1__Bacteroidetes;D_2__Bacteroidia;D_3__Chitinophagales;D_4__Saprospiraceae;D_5__Aureispira

D_0__Bacteria;D_1__Bacteroidetes;D_2__Bacteroidia;D_3__Chitinophagales;D_4__Saprospiraceae;__

D_0__Bacteria;D_1__Proteobacteria;D_2__Gammaproteobacteria;D_3__Pseudomonadales;D_4__Pseudomonadaceae;D_5__Pseudomonas

| | | | | |
|---|---|---|---|---|
| -1.257671573 | 13 | | 1.261889459 | 11 |
| -1.701843212 | 13 | | 1.706061098 | 11 |
| -1.296974807 | 12 | | 1.301192693 | 10 |
| 0.099951105 | 11 | | -0.095733219 | 9 |
| -0.19518464 | 11 | NA | | NA |
| -0.829507002 | 11 | | 0.833724888 | 9 |
| -1.45115789 | 11 | | 1.455375776 | 9 |
| 0.991162488 | 10 | | -0.986944602 | 9 |
| -0.827188412 | 10 | | 0.831406298 | 8 |
| -0.603899418 | 9 | | 0.608117304 | 7 |
| -0.410028872 | 8 | | 0.414246758 | 6 |
| -0.835911249 | 8 | | 0.840129135 | 6 |
| -0.748457653 | 5 | | 0.752675539 | 3 |

**Table S5:** Information on sample type, host population, diet, sampling time point, as well as number of raw ar

| sample id | individual | type | population | diet | day | time | raw reads |
|---|---|---|---|---|---|---|---|
| 2021_MB_1 | gosling1 | gut | gosling | brine shrimp | 7 | pm | 39602 |
| 2021_MB_2 | gosling2 | gut | gosling | brine shrimp | 7 | pm | 23079 |
| 2021_MB_3 | sayward1 | gut | sayward | brine shrimp | 12 | pm | 35054 |
| 2021_MB_4 | sayward2 | gut | sayward | brine shrimp | 11 | pm | 13397 |
| 2021_MB_5 | gosling3 | gut | gosling | brine shrimp | 7 | pm | 18099 |
| 2021_MB_6 | gosling4 | gut | gosling | brine shrimp | 12 | pm | 28704 |
| 2021_MB_7 | sayward3 | gut | sayward | brine shrimp | 10 | pm | 17837 |
| FECAL_2A | gosling1 | fecal | gosling | brine shrimp | 1 | pm | 65363 |
| FECAL_2B | gosling1 | fecal | gosling | brine shrimp | 2 | pm | 51359 |
| FECAL_2C | gosling1 | fecal | gosling | brine shrimp | 3 | pm | 86869 |
| FECAL_2D | gosling1 | fecal | gosling | brine shrimp | 5 | pm | 62371 |
| FECAL_2E | gosling1 | fecal | gosling | brine shrimp | 6 | pm | 73057 |
| FECAL_2F | gosling1 | fecal | gosling | brine shrimp | 7 | pm | 42018 |
| FECAL_3A | gosling2 | fecal | gosling | brine shrimp | 1 | pm | 46672 |
| FECAL_3B | gosling2 | fecal | gosling | brine shrimp | 2 | pm | 67772 |
| FECAL_3C | gosling2 | fecal | gosling | brine shrimp | 3 | pm | 50078 |
| FECAL_3D | gosling2 | fecal | gosling | brine shrimp | 4 | pm | 73502 |
| FECAL_3E | gosling2 | fecal | gosling | brine shrimp | 5 | pm | 66166 |
| FECAL_3F | gosling2 | fecal | gosling | brine shrimp | 7 | pm | 32963 |
| FECAL_4A | sayward1 | fecal | sayward | brine shrimp | 1 | pm | 23808 |
| FECAL_4B | sayward1 | fecal | sayward | brine shrimp | 3 | pm | 51191 |
| FECAL_4C | sayward1 | fecal | sayward | brine shrimp | 6 | pm | 37929 |
| FECAL_4D | sayward1 | fecal | sayward | brine shrimp | 8 | pm | 16905 |
| FECAL_4E | sayward1 | fecal | sayward | brine shrimp | 9 | pm | 44615 |
| FECAL_4F | sayward1 | fecal | sayward | brine shrimp | 12 | pm | 38797 |
| FECAL_6A | sayward2 | fecal | sayward | brine shrimp | 1 | pm | 52965 |
| FECAL_6B | sayward2 | fecal | sayward | brine shrimp | 2 | pm | 24257 |
| FECAL_6C | sayward2 | fecal | sayward | brine shrimp | 7 | pm | 48264 |
| FECAL_6D | sayward2 | fecal | sayward | brine shrimp | 8 | pm | 43335 |
| FECAL_6E | sayward2 | fecal | sayward | brine shrimp | 10 | pm | 52192 |
| FECAL_6F | sayward2 | fecal | sayward | brine shrimp | 11 | pm | 37022 |
| FECAL_7A | gosling4 | fecal | gosling | brine shrimp | 1 | pm | 29770 |
| FECAL_7B | gosling4 | fecal | gosling | brine shrimp | 3 | pm | 53438 |
| FECAL_7C | gosling4 | fecal | gosling | brine shrimp | 7 | pm | 31688 |
| FECAL_7D | gosling4 | fecal | gosling | brine shrimp | 10 | pm | 40362 |
| FECAL_7E | gosling4 | fecal | gosling | brine shrimp | 11 | pm | 39642 |
| FECAL_7F | gosling4 | fecal | gosling | brine shrimp | 12 | pm | 36319 |
| FECAL_8A | gosling3 | fecal | gosling | brine shrimp | 1 | pm | 17897 |
| FECAL_8B | gosling3 | fecal | gosling | brine shrimp | 2 | pm | 15136 |
| FECAL_8C | gosling3 | fecal | gosling | brine shrimp | 3 | pm | 54480 |
| FECAL_8D | gosling3 | fecal | gosling | brine shrimp | 5 | pm | 45063 |
| FECAL_8E | gosling3 | fecal | gosling | brine shrimp | 6 | pm | 40141 |
| FECAL_8F | gosling3 | fecal | gosling | brine shrimp | 7 | pm | 30853 |
| FECAL_9A | sayward3 | fecal | sayward | brine shrimp | 1 | pm | 24910 |
| FECAL_9B | sayward3 | fecal | sayward | brine shrimp | 2 | pm | 22533 |
| FECAL_9C | sayward3 | fecal | sayward | brine shrimp | 5 | pm | 47519 |
| FECAL_9D | sayward3 | fecal | sayward | brine shrimp | 6 | pm | 40680 |

| | | | | | | | |
|---|---|---|---|---|---|---|---|
| FECAL_9E | sayward3 | fecal | sayward | brine shrimp | 8 | pm | 19975 |
| FECAL_9F | sayward3 | fecal | sayward | brine shrimp | 10 | pm | 68654 |
| FECAL_11A | gosling5 | fecal | gosling | brine shrimp | 1 | pm | 81885 |
| FECAL_11B | gosling5 | fecal | gosling | brine shrimp | 2 | pm | 65505 |
| FECAL_11C | gosling5 | fecal | gosling | blood worms | 5 | am | 38374 |
| FECAL_11D | gosling5 | fecal | gosling | blood worms | 6 | am | 46618 |
| FECAL_11G | gosling5 | fecal | gosling | blood worms | 9 | am | 59896 |
| FECAL_11I | gosling5 | fecal | gosling | blood worms | 11 | am | 51869 |
| FECAL_11K | gosling5 | fecal | gosling | blood worms | 13 | am | 51266 |
| FECAL_12B | gosling6 | fecal | gosling | brine shrimp | 1 | pm | 47059 |
| FECAL_12C | gosling6 | fecal | gosling | brine shrimp | 2 | pm | 49042 |
| FECAL_12D | gosling6 | fecal | gosling | blood worms | 5 | pm | 53083 |
| FECAL_12E | gosling6 | fecal | gosling | blood worms | 7 | pm | 55367 |
| FECAL_12F | gosling6 | fecal | gosling | blood worms | 10 | am | 71764 |
| FECAL_12H | gosling6 | fecal | gosling | blood worms | 12 | am | 48671 |
| FECAL_12I | gosling6 | fecal | gosling | blood worms | 13 | am | 62988 |
| FECAL_13B | gosling7 | fecal | gosling | brine shrimp | 1 | pm | 78799 |
| FECAL_13C | gosling7 | fecal | gosling | brine shrimp | 2 | pm | 60162 |
| FECAL_13D | gosling7 | fecal | gosling | blood worms | 5 | am | 62195 |
| FECAL_13E | gosling7 | fecal | gosling | blood worms | 5 | pm | 57110 |
| FECAL_13F | gosling7 | fecal | gosling | blood worms | 6 | am | 58802 |
| FECAL_13G | gosling7 | fecal | gosling | blood worms | 6 | pm | 57291 |
| FECAL_13H | gosling7 | fecal | gosling | blood worms | 7 | am | 43782 |
| FECAL_13I | gosling7 | fecal | gosling | blood worms | 7 | pm | 43908 |
| FECAL_13J | gosling7 | fecal | gosling | blood worms | 8 | am | 45529 |
| FECAL_13K | gosling7 | fecal | gosling | blood worms | 8 | pm | 55932 |
| FECAL_13M | gosling7 | fecal | gosling | blood worms | 11 | am | 69526 |
| FECAL_13O | gosling7 | fecal | gosling | blood worms | 13 | am | 36793 |
| FECAL_14C | gosling8 | fecal | gosling | brine shrimp | 1 | pm | 62892 |
| FECAL_14D | gosling8 | fecal | gosling | brine shrimp | 2 | pm | 39239 |
| FECAL_14E | gosling8 | fecal | gosling | blood worms | 4 | pm | 47074 |
| FECAL_14F | gosling8 | fecal | gosling | blood worms | 5 | am | 55364 |
| FECAL_14I | gosling8 | fecal | gosling | blood worms | 8 | am | 46564 |
| FECAL_14K | gosling8 | fecal | gosling | blood worms | 11 | am | 50568 |
| FECAL_14M | gosling8 | fecal | gosling | blood worms | 13 | am | 50799 |
| FECAL_15C | gosling9 | fecal | gosling | brine shrimp | 1 | pm | 38136 |
| FECAL_15D | gosling9 | fecal | gosling | brine shrimp | 2 | pm | 31384 |
| FECAL_15F | gosling9 | fecal | gosling | blood worms | 5 | pm | 92728 |
| FECAL_15G | gosling9 | fecal | gosling | blood worms | 9 | am | 50547 |
| FECAL_15I | gosling9 | fecal | gosling | blood worms | 12 | am | 33543 |
| FECAL_15J | gosling9 | fecal | gosling | blood worms | 13 | am | 42337 |
| BLOODWORMS_1 | NA | diet | NA | NA | NA | NA | 37535 |
| BLOODWORMS_2 | NA | diet | NA | NA | NA | NA | 54709 |
| BRINESHRIMP_1 | NA | diet | NA | NA | NA | NA | 53758 |
| BRINESHRIMP_2 | NA | diet | NA | NA | NA | NA | 47597 |

nd filtered sequencing reads for all samples.

| filtered reads-no diet filtering | filtered reads-diet filtering |
|---|---|
| 11615 | 11178 |
| 9927 | 9706 |
| 7256 | 7147 |
| 7055 | 4738 |
| 5038 | 4238 |
| 12411 | 12411 |
| 11644 | 5448 |
| 37504 | 36060 |
| 38872 | 36928 |
| 51615 | 41531 |
| 42414 | 42007 |
| 48001 | 47636 |
| 23088 | 22550 |
| 30541 | 28578 |
| 40330 | 37448 |
| 35073 | 34067 |
| 51264 | 50712 |
| 45422 | 43594 |
| 21770 | 21731 |
| 8966 | 8212 |
| 41375 | 39772 |
| 32124 | 30279 |
| 15159 | 14486 |
| 37427 | 36818 |
| 34232 | 33406 |
| 46203 | 46016 |
| 19236 | 18060 |
| 39566 | 38476 |
| 32235 | 30944 |
| 32845 | 32398 |
| 31306 | 30544 |
| 20788 | 19926 |
| 37604 | 37420 |
| 23311 | 22608 |
| 29668 | 28975 |
| 24064 | 23414 |
| 30213 | 30050 |
| 12164 | 12086 |
| 7391 | 7308 |
| 41186 | 39715 |
| 29662 | 29098 |
| 26590 | 25634 |
| 22136 | 20871 |
| 23111 | 23084 |
| 19950 | 18830 |
| 37160 | 36772 |
| 34851 | 34402 |

| | |
|---|---|
| 16283 | 16051 |
| 61037 | 60861 |
| 38968 | 16925 |
| 45003 | 40175 |
| 28125 | 19673 |
| 32289 | 12835 |
| 42765 | 17382 |
| 38184 | 12341 |
| 36835 | 13205 |
| 29102 | 11779 |
| 23998 | 18194 |
| 40919 | 36437 |
| 43223 | 26617 |
| 37198 | 19491 |
| 38571 | 21103 |
| 43234 | 36478 |
| 45889 | 29275 |
| 22460 | 18789 |
| 43695 | 15420 |
| 41675 | 23456 |
| 43039 | 12663 |
| 40175 | 27812 |
| 33592 | 12954 |
| 28881 | 14089 |
| 33218 | 7560 |
| 15432 | 4275 |
| 51357 | 28979 |
| 26244 | 12153 |
| 46987 | 7801 |
| 24681 | 16913 |
| 19051 | 12246 |
| 42505 | 25174 |
| 32291 | 13378 |
| 38328 | 19248 |
| 34999 | 24390 |
| 20127 | 8931 |
| 11182 | 8416 |
| 67901 | 29522 |
| 42965 | 19615 |
| 26730 | 11168 |
| 31917 | 17748 |
| 31863 | NA |
| 47231 | NA |
| 42569 | NA |
| 39315 | NA |
